# Characterization of apo-form selective inhibition of indoleamine 2,3-dioxygenase

**DOI:** 10.1101/324947

**Authors:** Rodrigo Ortiz-Meoz, Liping Wang, Rosalie Matico, Anna Rutkowska, Martha De la Rosa, Sabrina Bedard, Robert Midgett, Katrin Strohmer, Douglas Thomson, Cunyu Zhang, Makda Mebrahtu, Jeffrey Guss, Rachel Totoritis, Thomas Consler, Nino Campobasso, David Taylor, Tia Lewis, Kurt Weaver, Marcel Mülbaier, John Seal, Richard Dunham, Wieslaw Kazmierski, David Favre, Giovanna Bergamini, Lisa Shewchuk, Alan Rendina, Guofeng Zhang

**Affiliations:** Drug Design and Selection, GlaxoSmithKline, 1250 S. Collegeville Rd, Collegeville, Pennsylvania, 19426, United States; Cellzome GmbH, GlaxoSmithKline, Meyerhofstrasse 1, D-69117 Heidelberg, Germany; Infectious Diseases TAU, GlaxoSmithKline, Five Moore Drive, Research Triangle Park, North Carolina, 27709, United States

## Abstract

Indoleamine-2,3-dioxygenase 1 (IDO1) is a heme-containing enzyme that catalyzes the rate-limiting step in the kynurenine pathway of tryptophan (TRP) metabolism. As an inflammation-induced immunoregulatory enzyme, pharmacological inhibition of IDO1 activity is currently being pursued as a potential therapeutic tool for the treatment of cancer and other disease states. As such, a detailed understanding of the mechanism of action of established and novel IDO1 inhibitors remains of great interest. Comparison of a newly-developed IDO1 inhibitor (GSK5628) to the existing best-in-class compound, epacadostat (Incyte), allows us to report on a unique inhibition mechanism for IDO1. Here, we demonstrate that GSK5628 inhibits IDO1 by competing with heme for binding to a heme-free conformation of the enzyme (apo-IDO1) while epacadostat coordinates its binding with the iron atom of the IDO1 heme cofactor. Comparison of these two compounds in cellular systems reveals a long-lasting inhibitory effect of GSK5628, undescribed for other known IDO1 inhibitors. Detailed characterization of this apo-binding mechanism for IDO1 inhibition may help design superior inhibitors or may confer a unique competitive advantage over other IDO1 inhibitors *vis-à-vis* specificity and pharmacokinetic parameters.

## INTRODUCTION

In mammals, indoleamine-2,3-dioxygenase (IDO1) expression and activity is induced during inflammation, initiating the kynurenine pathway of tryptophan catabolism and promoting an immunosuppressive local environment.^1–4^ This immunosuppression is suggested to prevent rejection of the mammalian fetus by the maternal immune system by suppressing T cell activity and may play a role in limiting tissue damage by an overexuberant immune response in other settings.^5–6^ As the first catalytic step in tryptophan (TRP) metabolism, upon expression, IDO1 acts on TRP to produce N-formylkynurenine (NFK). Subsequent steps in the kynurenine pathway of TRP metabolism produce several metabolites that are thought to play immunosuppressive roles, including kynurenine (KYN) and 3-hydroxyanthranilic acid (3-HAA). KYN is anti-proliferative and pro-apoptotic to T cells and NK cells responding to antigenic stimuli while 3-HAA inhibits macrophage function and promotes the differentiation of immunosuppressive regulatory T cells (Treg). ^7–9^

A number of disease states are associated with IDO1 induction and immunosuppressive function. IDO1 expression and activity are induced during untreated HIV infection and remain elevated despite antiretroviral therapy (ART) suppression of virus replication.^10–11^ Increased IDO1 expression levels correspond with the risk of non-AIDS morbidity/mortality in several cohorts of ART-suppressed HIV-infected patients.^12–14^ In Oncology, IDO1 is expressed by tumor cells in multiple tumor types; some studies finding expression of IDO1 in 58% of tumor samples tested.^2^ As with HIV patients, IDO1 expression in cancerous cells has also been linked with poor patient prognosis and increased tumor progression.^15^ The expression of IDO1 in human tumors and the resulting ability of the tumor to escape immune response has been the subject of many studies ^16–19^ and a significant investment in the discovery of potent, selective IDO1 inhibitors.^20–21^ To date, numerous clinical trials involving IDO1 inhibition alone or in combination are being pursued.^22^ The recent termination of a phase III clinical trial (ECHO-301 / KEYNOTE-252) which combined epacadostat (**Figure 1**), an IDO1 inhibitor developed by Incyte, with Merck’s pembromuzilab has precipitated the clinical need for an alternative mechanism of inhibiting IDO1. IDO1 is a heme-containing, monomeric oxygenase that catalyzes the di-oxygenation of tryptophan to form NFK. As is common with other inhibitors that target metalloproteins, epacadostat coordinates with the catalytic metal, in this case a heme iron, via its N-hydroxyamidine moeity.^23–26^ Coordination with the heme-sequestered iron in conjunction with built-in affinity to the IDO1 tryptophan binding pocket is a mechanism of inhibition shared by several reported IDO1 inhibitors.^27^ Here, we report detailed studies of the mechanism of action of GSK5628 (**Figure 1**), a potent and selective IDO1 inhibitor that does not coordinate with the heme iron via a metal binding group but instead competes with heme for binding to a heme-free form of IDO1 (apo-IDO1), a novel, recently described inhibition mechanism.^28^

**Figure 1.**
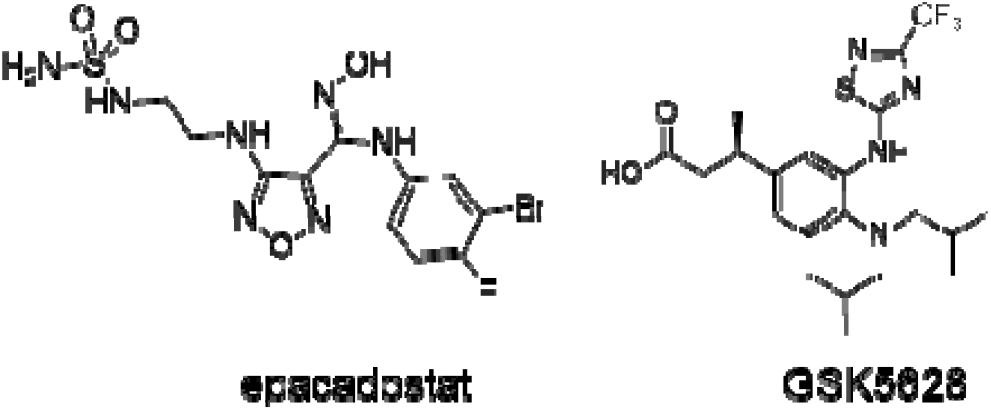
epacadostat and GSK5628 inhibit IDO1

## RESULTS AND DISCUSSION

### GSK5628 Inhibits IDO1 in Cells

Initially, to assess the potency of GSK5628, we employed a cell-based assay where IDO1 expression is induced with human interferon-gamma (IFNγ) in HeLa cells (see SI Methods).^29^ IDO1 expression and activity results in the conversion of TRP to NFK by IDO1 and the subsequent processing of NFK to kynurenine (KYN) by endogenous NFK formamidase. In this assay, which measures the accumulation of KYN in the cell media, epacadosta and GSK5628 both exhibit robust IDO1 inhibition (**24 nM and 5.9 nM IC_50_, respectively**, **Figure S1A**) without causing cell cytotoxicity as measured by CellTiter-Glo (Promega) (**Figure S1B**). These results agree with previously published reports of epacadostat and show that GSK5628 is a potent inhibitor of IDO1 in a cellular context.^21^

### Increased Time and Temperature Facilitates GSK5628 Inhibition *in vitro*

Measuring the potency of GSK5628 in a biochemical assay in which recombinant IDO1 activity is measured by following the production of NFK from a D-TRP substrate (see SI Methods) yielded surprising results compared to epacadostat.^21, 30^ When this biochemical assay is run at 25°C and with a 15-minute Enzyme: Inhibitor preincubation before addition of the substrate, epacadostat inhibits IDO1 (81 nM IC_50_) but, paradoxically, GSK5628 does not exhibit any significant inhibition of IDO1 (**Figure 2A**). In light of these results, and in order to determine whether the robust inhibition of KYN formation by GSK5628 seen in the cellular assay was due to bona fide IDO1 target engagement, we performed a cellular thermal shift assay (CETSA)^31–32^ in HeLa cells. Results from this assay show that GSK5628 can effectively bind and thermally stabilize IDO1 in cells (**Figure S2**), confirming that GSK5628 engages IDO1 in a cellular context. To understand the disconnect between the cellular potency of GSK5628 and inactivity in the recombinant-IDO1 biochemical assay, many of the parameters of the assay were varied, including preincubation duration and temperature. We observed that increased temperature from 25°C to 37°C as well as increased duration of Enzyme: Inhibitor preincubation (before addition of substrate) from 0 to 135 minutes led to increased inhibition of IDO1 by GSK5628 in the biochemical assay. Full inhibition was achieved with a 120-min incubation at 37°C (**Figure S3**). Thus, while increasing the temperature to 37°C and the pre-incubation time to 120 min caused no effect on the calculated potency of epacadostat (82 nM versus 62 nM IC_50_), this change had a profound effect on GSK5628 (**Figure 2B**), in essence “rescuing” the activity of the compound (little to no activity to 39 nM IC_50_) and resolving the original discrepancy between the cellular and biochemical potency of GSK5628. Fitting this data to the Morrison tight binding equation returns Ki values of 29 nM for epacadostat at 25°C, 20 nM for epacadostat at 37°C and 10 nM for GSK5628 at 37°C (**Figure S4**). Follow-up studies concluded that a 120 min / 37°C incubation of IDO1 in the absence of GSK5628 followed by addition of GSK5628 did not result in increased inhibition (**Figure S5**) and that, at 25°C, the activity of GSK5628 can be observed, but inhibition increases at a much slower time scale (**Figure 4B**).

**Figure 2.**
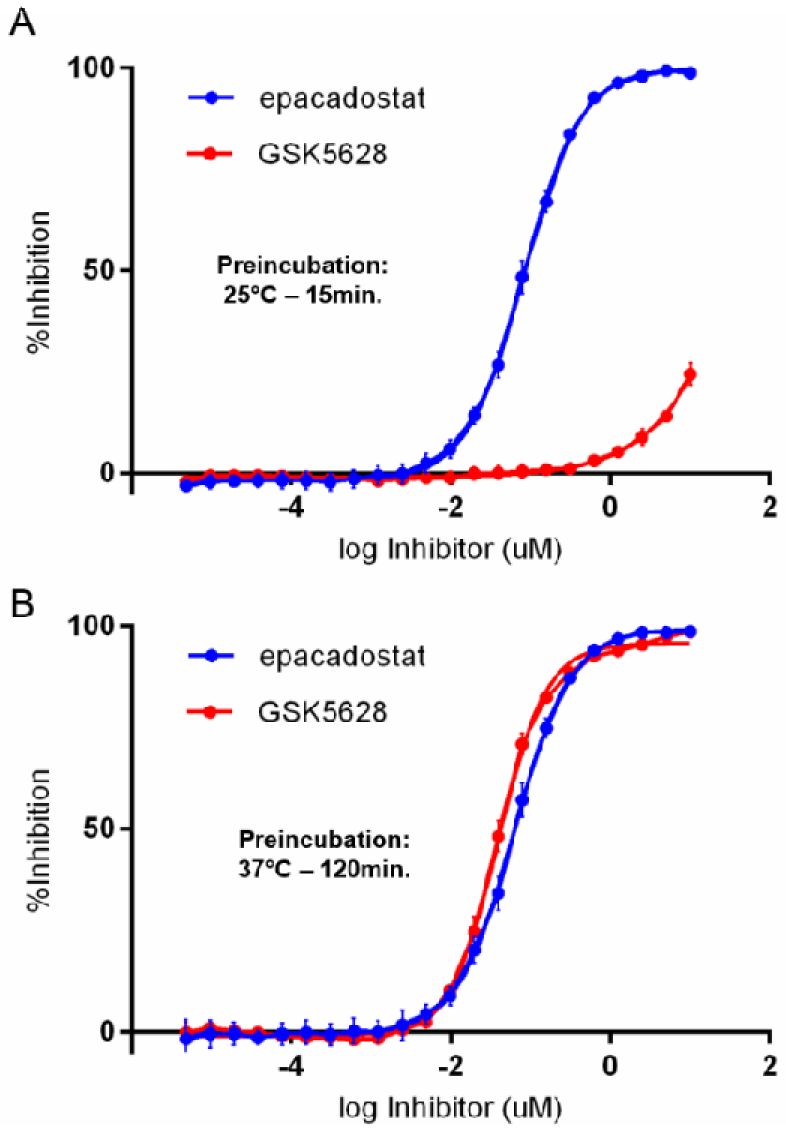
GSK5628 requires preincubation with enzyme to inhibit IDO1. (A) epacadostat (blue line) but not GSK5628 (red line) inhibits IDO1 in a biochemical assay with a 15 min / 25°C enzyme: inhibitor preincubation. Increasing the preincubation time and temperature to 120 min / 37°C (B) rescues the activity of GSK5628 (red line) while not affecting epacadostat (blue line) potency. All data is average of n=4 replicates with the standard deviation.

### GSK5628A Competes with the IDO1 Heme Cofactor

As a heme-containing protein, IDO1 has a characteristic absorption peak at 405 nm. This Soret band is known to report on the electronic state of the iron in heme-containing proteins ^33–34^ and, as expected, IDO1 incubated (37°C / 120 min) with epacadostat causes a red-shift in the Soret band indicative of coordination of the heme iron^26^ (**Figure 3A**). Surprisingly, incubation of IDO1 and GSK5628 at 37°C for 120 minutes leads to ablation of the 405 nm IDO1/heme peak and appearance of a new peak at a shorter wavelength similar to that of free heme (**Figure 3A**). This ablation of the Soret band was not observed when IDO1 was incubated with inactive analogs of GSK5628 (data not shown). To understand whether this phenomenon is specific to IDO1, several additional non-covalent heme-containing proteins were studied, including human hemoglobin, myoglobin, and tryptophan 2,3-dioxygenase (TDO2). There was no observed loss of heme and reduction of the Soret band in these proteins after incubation with GSK5628 at 37°C for 120 minutes (**Figure S6**). These observations suggest that the incubation of GSK5628 and IDO1 leads to the loss of heme, and the loss of heme is specific and selective. To further assess the lo s of heme in the presence of GSK5628, we measured the free heme after incubation with inhibitor using an apo-peroxidase assay for free heme detection (SIGMA MAK-036). We observed that, after a 37°C /120-m nute incubation, epacadostat decreases the amount of free heme relative to a DMSO-control while GSK5628 causes a significant increase in free heme relative to DMSO and epacadostat (**Figure 3B**). The epacadostat-mediated decrease is consistent with the fact that epacadostat coordinates its binding with the heme iron and this interaction presumably helps stabilize heme in the IDO1 pocket. One explanation for the increased free heme upon GSK5628 treatment is that GSK5628 displaces heme from IDO1 as it binds, and suggests that GSK5628 and the heme cofactor compete for the ame site. To understand which region of IDO1 is affected by GSK5628 binding, hydrogen-deuterium exchange mass spectrometry (HDX-MS^35^) was performed at 37°C with a 120 min preincubation.^36^ This result **(Figure 3C**, **S7A and S7B**) shows that GSK5628 protects residues around the heme-binding pocket of IDO1, indicating that GSK5628 and the IDO1 heme cofactor both bind in the IDO1 active site.

**Figure 3.**
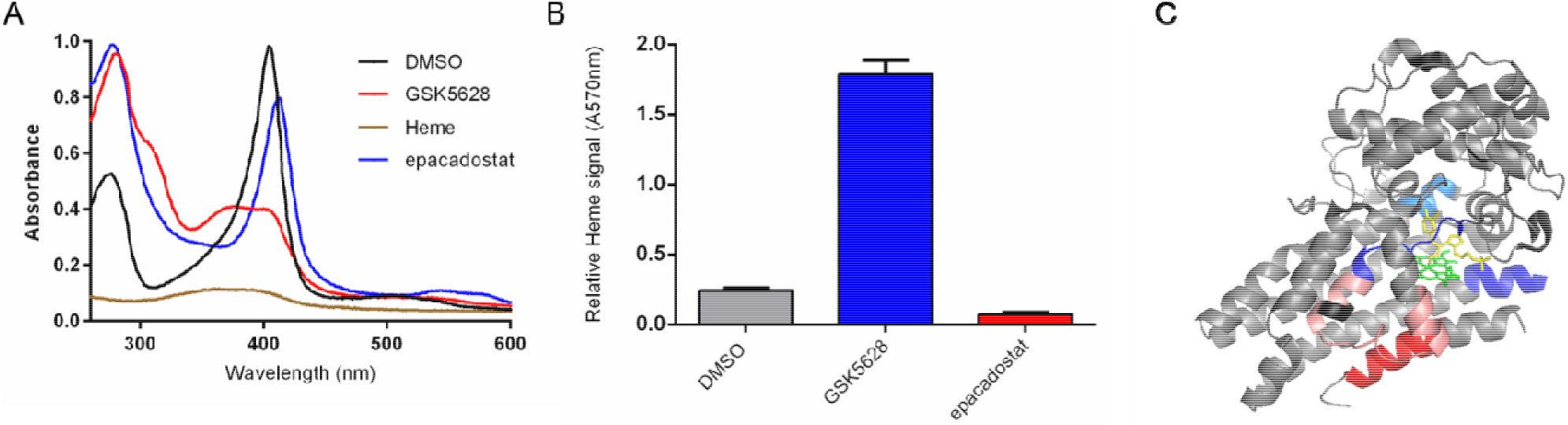
GSK5628 and heme compete for the same binding site. (A) Epacadostat (blue line) shifts the Soret band of IDO1 after binding but GSK5628 binding (red line) results in a loss of this band, with a new spectrum similar to unincorporated heme (brown line). (B) GSK5628 incubation with IDO1 causes a significant increase in free heme (red bar) while epacadostat protects heme release from IDO1 (blue bar). (C) Differences in hydrogen-deuterium exchange, between heme-bound and GSK5628-bound IDO1, mapped onto the epacadostat-bound IDO1 structure (PBD ID 5WN8^26^) show that GSK5628 binding protects residues (blue) around the heme (green) and epacadostat (yellow) binding sites. Red residues show areas where GSK5628 causes deprotection. Residues that do not show a significant difference in exchange between heme-bound and GSK5628-bound are colored in gray while residues with no exchange information are in black.

### GSK5628 Binds Apo-IDO1

The shared binding site of GSK5628 and heme, combined with the observation that GSK5628 gradually inhibits IDO1 as the enzyme and compound are incubated at 37°C, led us to propose a novel mechanism of inhibition for GSK5628 (**Figure 4A**).^28^ Central to this model is the observation that IDO1 can freely exist in equilibrium between its heme-bound (**holo-IDO1**) and apo (**apo-IDO1**) forms. As a non-covalently bound cofactor, the IDO1 heme necessarily exists in a binding equilibrium and, not surprisingly, mutating resides around the IDO1 heme-binding site seems to perturb this equilibrium.^37^ In this model, epacadostat acts as a classic metalloenzyme inhibitor; coordinating the heme iron in holo-IDO1 while GSK5628 competes with heme and exclusively binds to the apo form of IDO1. As molecules of GSK5628 bind apo-IDO1, they prevent the reloading of the heme cofactor, effectively inhibiting reconstitution of active IDO1 (holo-IDO1). The competition of GSK5628 and heme for the same IDO1 site helps to explain the required preincubation time for GSK5628 to inhibit a starting enzyme population of nearly 100% holo-IDO1, where the dissociation of heme from holo-IDO1 is the rate-limiting step. A similar mechanism was proposed for 3 IDO1 inhibitors described in Nelp et al.^28^, work which culminated in crystal structures of these inhibitors bound to apo-IDO1 (PDB ID 6AZV). As expected, the HDX-MS differences we observe for GSK5628 mapped onto this apo-IDO1 structure (**Figure S7C)** show that GSK5628 protects residues around the heme-binding pocket of IDO1. Furthermore, a docking model (**Figure S8A**) shows that GSK5628 has significant overlap with BMS-978587, suggesting that these molecules share the same mechanism of inhibition.^28^

**Figure 4.**
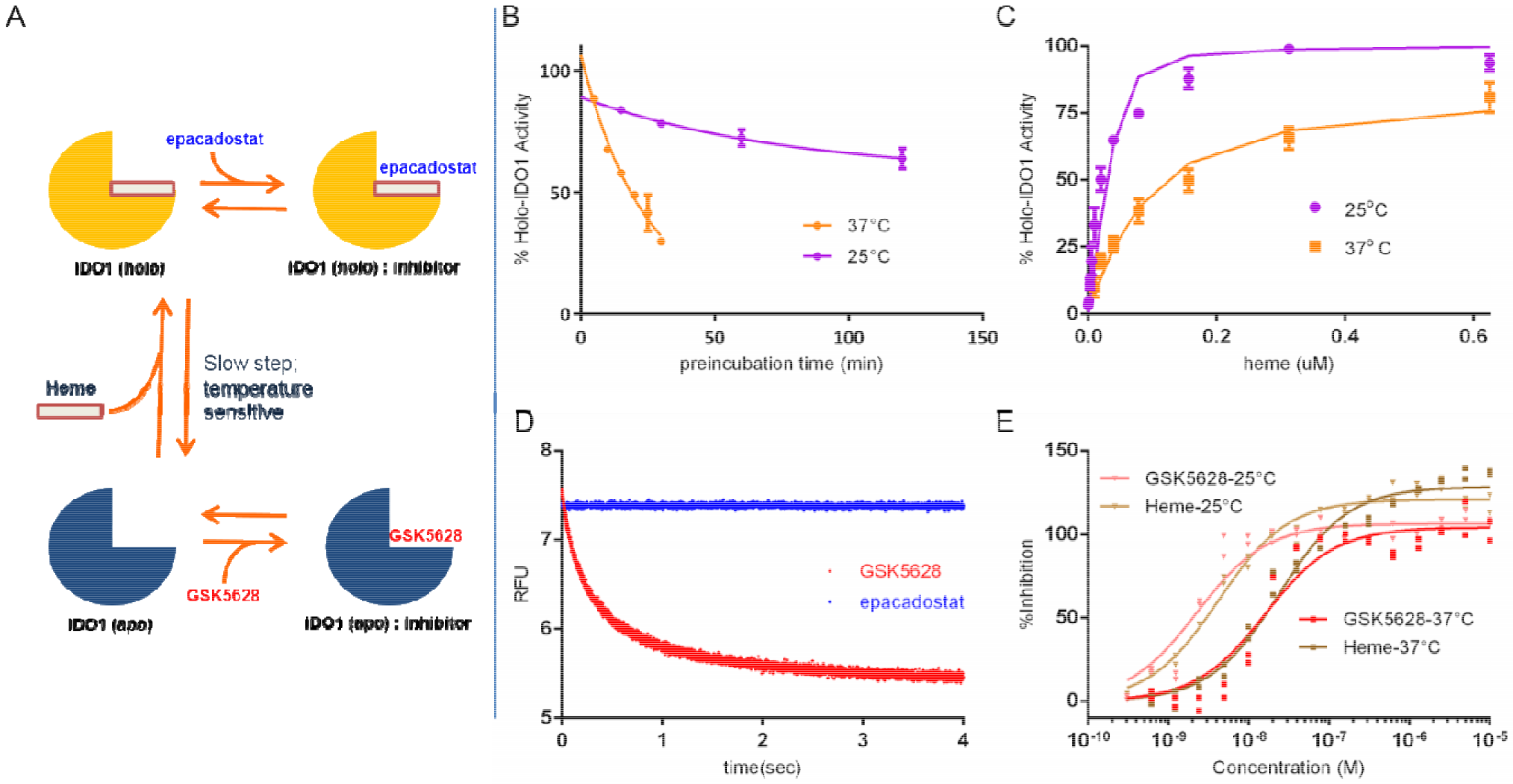
GSK5628 binds apo-IDO1. (A) Scheme of equilibrium between holo-IDO1 and apo-IDO1 depicting different binding modes of epacadostat and GSK5628. (B) holo-IDO1 (50 nM) was incubated with 5 uM GSK5628 for the indicated times at 25°C (purple line) and 37°C (orange line) and then assayed for activity. (C) apo-IDO1 can bind heme and reconstitute catalytic activity. Measurements performed at 25°C(purple line) and 37°C (orange line) (D) GSK5628 (1 mM; red line) can bind directly and rapidly to 250 nM apo-IDO1 and quench the intrinsic TRP fluorescence, but epacadostat (1 mM; blue line) cannot. (E) Equilibrium competitive binding studies using apo-IDO1 and a fluorescent probe were performed at 25°C (triangles) and 37°C (circles), highlighting the change in heme and GSK5628 IC_50_ with a change in temperature. Potency is underestimated due to the relatively high apo-IDO1 / probe ratio needed for the FP assay format.

A key assumption in this model is the ability of IDO1 to exist stably in the apo form and to be capable of binding heme and reconstituting its dioxygenase activity. Purification of apo-IDO1 is described in detail in the Methods and a full characterization of the apo protein can be found in **Figure S9** where we show that apo IDO1 is stable and not prone to aggregation under the conditions used in this study, consistent with other heme-containing proteins, which can exist as stable apo forms ^38–42^. To test whether this apo-IDO1 preparation can be returned to the holo form, the catalytic activity of recombinant apo-IDO1 was measured against an increasing amount of free heme (**Figure 4C**). Without addition of free heme, apo-IDO1 has no catalytic activity and as the heme concentration is increased, the catalytic activity is returned, reaching an overall level similar to a holo-IDO1 control (100% activity). Assuming that the fraction of heme-bound IDO1 can be extrapolated from the catalytic activity of the enzyme, this experiment also allows us to estimate the affinity of apo-IDO1 for heme at 25°C (4.6 nM) and 37°C (59.7 nM) using a tight binding substrate equation (See **SI Methods**). The dissociation rate of heme from holo-IDO1, which we predict to be the slow, temperature sensitive step that necessitates the 120 min / 37°C pre-incubation for GSK5628 to achieve maximum potency, can be estimated by extrapolating the inhibition data of a saturating amount of GSK5628 as a function of time at both 25°C and 37°C (**Figure 4B**). These results show that the heme apparent off rate constant from holo-IDO1 is 0.0007 s^−1^ (t_1/2off_ = 16.4 min) at 37 °C and 0.0002 s^−1^ (t_1/2off_ = 54.5 min) at 25 °C, consistent with the proposed model with the heme off rate being the rate-limiting step in GSK5628 inhibition.^43^ Next, we measured the direct binding of GSK5628 and epacadostat to apo-IDO1 by following the intrinsic fluorescence of the protein (**Figures 4D and S10**). Here, we observe that while epacadostat is unable to bind to apo-IDO1, GSK5628 exhibits biphasic binding to the apo-form of IDO1 with k_on_ measurements of 1.2 and 0.056 μM^−1^s^−1^ at 25 °C and 3.7 and 0.65 μM^−1^s^−1^ at 37 °C for the fast and slow phases, respectively. These results show that, in the absence of heme, GSK5628 binds rapidly to IDO1 regardless of temperature. Furthermore, equilibrium titrations of the quenching of the intrinsic TRP fluorescence of apo-IDO1 provide accurate measures of the affinity at both temperatures (Kd = 2.1 nM at 25 °C and 5.0 nM at 37 °C, **Figure S11**). Affinity-Selection / Mass Spectrometry studies (AS/MS)^44–45^ show that GSK5628 binds to apo-IDO1 regardless of the duration or temperature of incubation and binding to holo-IDO1 is increased only as the time and temperature are increased (**Figure S12A**). Further, increasing the amount of heme present in the reaction diminishes the signal of the small amount of GSK5628 that can bind holo-IDO1 and is consistent with shifting the equilibrium of IDO1 in solution towards an increased holo population, to which GSK5628 cannot bind (**Figure S12B**).

### Fluorescent Probe Studies

To further explore the competition of GSK5628 and heme for the same binding pocket in IDO1, a fluorescent analog of GSK5628 was synthesized (**GSK1051**, **Figure S13A**). A docking model with GSK5628 shows the solvent exposed region that can accommodate fluorophores and click probes (**Figure S8B**). This fluorescent analog binds apo-IDO1 with a Kd of 10.6 nM at 25°C and 24.8 nM at 37°C (**Figure S13B and C**) and relatively rapid on and off rate constants (**Figure S14**), consistent with GSK1051 being a weaker derivative of GSK5628. Competition experiments (**Figure 4E**) following the change in fluorescence polarization (FP) of an apo-IDO1:GSK1051 complex show that both GSK5628 and heme can displace the probe at 25°C with high affinity (GSK5628 IC_50_: 2.2 nM, heme IC_50_: 4.4 nM). These values for the heme and GSK5628 IC_50_’s at 25 °C are consistent with the values we had previously calculated in two separate experiments (**Figure 4C and S11**). Repeating this experiment at 37°C (**Figure 4E**) shifts the measured IC_50_ of heme for apo-IDO1 to 27 nM, consistent with the extrapolated off rates at varying temperatures (**Figure 4B**) and explaining the temperature dependence of GSK5628 inhibition of IDO1. The IC_50_ of 14 nM for GSK5628 at 37°C is also consistent with the Kd value obtained by equilibrium titration of the intrinsic fluorescence (**Figure S11**). The accelerated dissociation of heme (and both GSK5628 and GSK1051) at 37°C could also be measured by kinetic studies of the ligand competition. Time courses at both 25°C and 37°C for competition with or direct binding of the fluorescent probe at multiple heme and GSK5628 and GSK1051 concentrations were fit globally using KinTek Explorer ^46–47^ software (**Figures S14, S15 and S16**). The k_off_ values for GSK5628 and GSK1051 increased ~4-5-fold from 0.00015 and 0.0088 s^−1^ at 25°C, respectively, to 0.00064 and 0.045 s^−1^ at 37°C. The k_off_ value for the heme increased ~3-fold from 0.00027 s^−1^ (t_1/2 off_ = 44 min) at 25 °C to 0.00073 s^−1^ (t_1/2 off_ = 16 min) at 37°C consistent with the earlier estimates and confirming that GSK5628 can effectively trap the apo form and that the slow temperature sensitive step in GSK5628 inhibition is heme dissociation.

### The Inhibitory Mechanism of GSK5628 in Cells

The aforementioned apo-binding mechanism of GSK5628 is predicated on the competition of heme and the molecule for the same binding site on the IDO1 peptide. To test whether this competition is also present in a cellular context, HeLa cells were incubated in the presence of 40 uM exogenous hemin and varying concentrations of GSK5628 and epacadostat. After incubation, the cellular potencies of the molecules were measured (**Figure S1C**). Consistent with the apo-binding mechanism, addition of exogenous heme shifts the potency of GSK5628 as competition prevents effective binding of the inhibitor while the potency of epacadostat remains unaltered. To assess the long-term consequences of the apo-binding mechanism of inhibition in a cellular context, we designed an experiment to measure the persistence of IDO1 inhibition after treatment with either epacadostat or GSK5628 in IFNγ-stimulated HeLa cells (**Figure S17**). As expected, a 24-hour incubation of IFNγ-stimulated HeLa cells with either epacadostat or GSK5628 results in the complete inhibition of IDO1 production as measured by KYN levels in the media (**Pre-washout**: **Figure 5A**). Subsequently, to test the reversibility of this inhibition, cells were washed with cycloheximide-containing media to ensure removal of the inhibitor and to prevent any new IDO1 synthesis. After another 22-hour incubation to allow KYN levels to replenish, KYN amounts were measured for cells that had been previously treated with GSK5628, epacadostat or DMSO. Consistent with our model, epacadostat-treated cells completely regain IDO1 activity after compound washout (**Figure 5A**). Surprisingly, GSK5628-treated cells do not show an activity increase commensurate with epacadostat or DMSO-treated cells (**Figure 5A**). Western blot analysis shows that this prolonged inhibitory effect of IDO1 activity is not due to a degradation of IDO1 upon GSK5628 treatment (**Figure S17**). In fact, GSK5628 treatment seems to lead to a slight increase in IDO1 protein levels, a commonly observed feedback mechanism upon small molecule inhibition of other protein targets ^48^ and previously observed in IDO1.^49^ Thus, it would appear that the population of cellular IDO1 existing in the cells after GSK5628 treatment and washout is not able to form an active complex with heme. To further investigate these findings, we developed a cell-permeable probe (**GSK5112**, **Figure S18**) based on GSK5628 that features a trans-cyclooctene (TCO) group to enable bioorthogonal click chemistry reaction with reporter dye and subsequent intracellular imaging. ^50–51^ Incubation of naïve and IFNγ-stimulated HeLa cells with GSK5112 followed by fixing and conjugation with a Cy5-tetrazine reporter indicates that GSK5112 is a selective probe as it can be detected only in the conditions where IDO1 expression has been induced (IFN-γ stimulation) (**Figure S18**). Predictably, both GSK5628 and epacadostat pretreatment inhibit probe binding to IDO1 (**Figure 5B**). In this case, GSK5628 effectively competes with the probe for apo-IDO1 sites while epacadostat stabilizes heme in holo-IDO1 and prevents the probe from competing with heme, thereby decreasing its binding. Using this probe, we can also interrogate the heme-associated state of the IDO1 population in HeLa cells after GSK5628 inhibition and washout with cycloheximide-containing media, conditions where we have shown that IDO1 activity remains absent (**Figure 5A**). After washout of the initial GSK5628A treatment, cells were incubated with either DMSO, GSK5628, or epacadostat 3 hours prior to addition of the probe (**Figure S19B**). If, after GSK5628 wash-out, heme would be able to re-bind IDO1, epacadostat should be able to inhibit GSK5112 binding, consistent with previous results (**Figure 5B)**. However, under these conditions, only GSK5628 but not epacadostat, was able to block GSK5112 binding (**Figure 5C**) indicating that the IDO1 pool present in cells after the washout consists mainly of apo-IDO1. Thus, the likely explanation of the durable inhibition observed with GSK5628 is indeed the limitation in reloading of heme.

**Figure 5.**
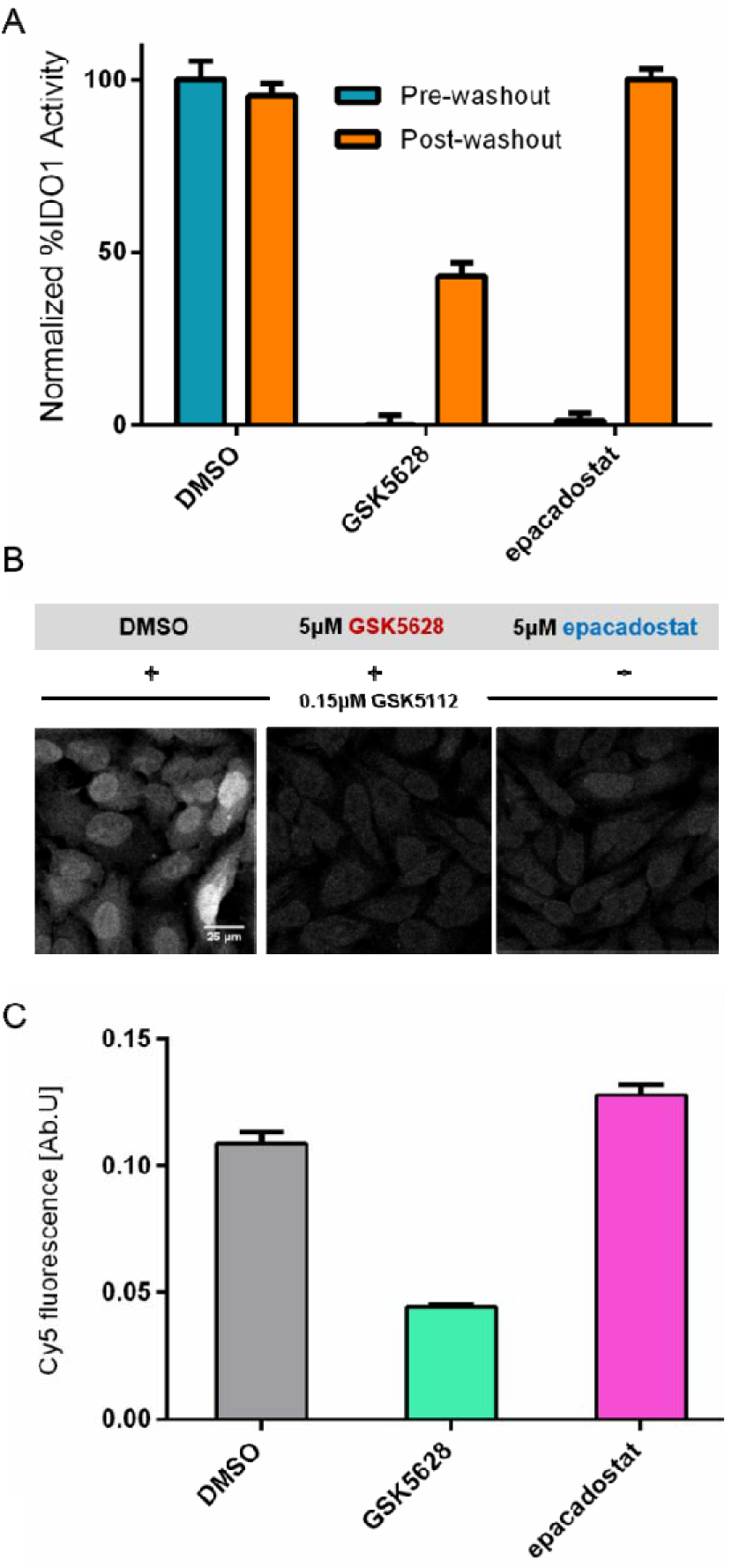
Prolonged inhibitory effect of GSK5628 in cells. (A) Washout experiments showing that GSK5628 but not epacadostat shows a durable decrease in IDO1 activity after a compound washout (orange bars). (B) GSK5112 engages IDO1 in the presence of DMSO but not epacadostat and GSK5628. (C) epacadostat (pink bar) does not block GSK5112 binding in post-washout GSK5628-treated cells.

## DISCUSSION

Here, through biochemical, biophysical, and cellular studies, we have shown that GSK5628 can effectively bind and inhibit IDO1 biochemically and in a cellular context. As we have shown, GSK5628 binds to the heme-free form of IDO1 (apo-IDO1). By competing with heme for the same site, this binding event prevents apo-IDO1 from forming an active complex with its cofactor, ensuring inhibition of catalytic activity. While we see a slow onset of inhibition for GSK5628 *in vitro* when compared to a more traditional compound which binds the active form of IDO1, we observe a durable inhibitory effect even after washout of GSK5628 in a cellular context. We propose that this durability is due to the unavailability of the heme cofactor in the cellular matrix. A fluorescent probe is able to readily engage this IDO1 pool, implying that it is properly folded and able to bind the heme cofactor but instead remains in the inactive apo form. During the preparation of this manuscript, Groves and coworkers also described the binding of IDO1 inhibitors to the apo form.^28^ Where our studies have overlapping, comparable data, (such as the calculation of the heme dissociation values at 37°C) we see significant agreement, strengthening the validity of the mechanism of inhibition described here. More recently, a study features a crystal structure of BMS-986205 (one of the inhibitors used by Nelp et al.^28^) bound initially to holo-IDO1 followed by several conformational changes that result in heme release and a final apo-IDO1-inhibitor complex^52^. While these results appear to contradict our findings and those of Nelp et al., the crystallographic and spectroscopic data supporting their conclusions were conducted at high protein and inhibitor concentrations suggesting that this is a low affinity pathway compared to the rapid, potent and direct interaction of these types of inhibitors with apo-IDO1 (**Figure S20** confirms potent, rapid and direct binding of BMS-986205 to apo-IDO1). Furthermore, their mechanism does not account for the reversible nature of inhibitor binding to apo-IDO1 clearly demonstrated here and in Nelp et al., nor does it account for the remarkably similar time and temperature dependence of holo-IDO1 inhibition shared by apo-myoglobin^28^, BMS-978587^28^, BMS-986205^28^ and GSK5628 (37°C t_1/2_ = 17, 14, 11 and 16 min, respectively, which match the heme dissociation measured here with apo-IDO1 (37°C t_1/2_ =16 min). We acknowledge that BMS-986205 could bind directly to holo-IDO1 under non-physiological conditions (**Figure S21** scheme), but the more relevant pathway in cells is to trap the apo-IDO1. These findings have broad implications for the potential physiological advantage of inhibiting apo-IDO1 in a clinical context and, more broadly, the role of heme biosynthesis / binding on the activity and regulation of IDO1. Further, this new mechanism of inhibition can conceivably be exploited to target the apo-form of enzymes with a non-covalently bound cofactor and could open up new avenues to interrogate previously untractable protein targets.

## ASSOCIATED CONTENT

### Supporting Information

Detailed experimental procedures, reagent generation and supporting figures. This material is available free of charge via the Internet at http://pubs.acs.org.

## AUTHOR INFORMATION

### Notes

The authors declare the following competing financial interest(s): Current and former employees of GlaxoSmithKline, plc.

## ACKNOWLEDGMENT

The authors would like to thank Shu Zhang for statistical analysis support. Eric Shi, Jonathan Basilla, Jessica Schneck, Geoffrey Quinque, Caterina Musetti and Jeffrey Gross for helpful discussions and suggestions during the experimental design and manuscript preparation.

---

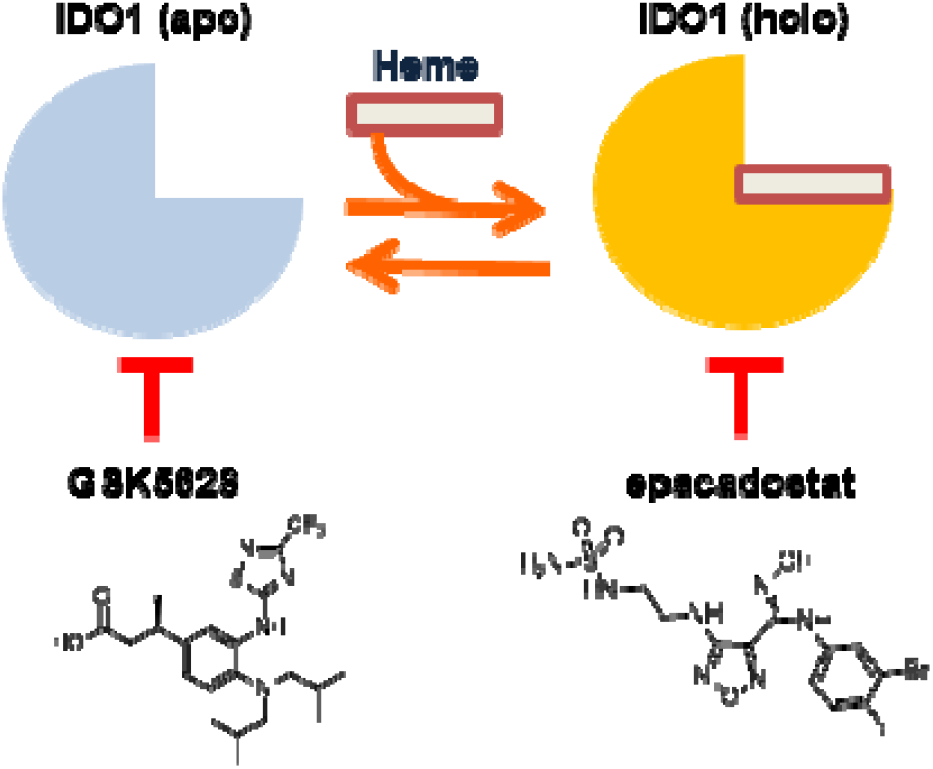

