## Supplementary Information for "Characterization of apo-form selective inhibition of indoleamine 2,3-dioxygenase"

**Supporting Information**

**Inhibitor Synthesis:**

**General Information**

Preparative high performance liquid chromatography (prep. HPLC) was performed on a Waters auto purification system [2767 Sample Manager, 2545 Binary Gradient Module, 2420 ELS Detector, 2996 Photodiode Array Detector, 3100 Mass Detector, Waters SFO, Waters 515 HPLC Pump]. The purification was done on a XBridge BEH C18 OBD 5µm Prep Column using a water/acetonitrile mixture (0.2% formic acid as modifier) at a flow rate of 30 mL/min. The purity of all final compounds was 95% or higher as determined by ultra performance liquid chromatography (UPLC MS), photodiode array detection. The instrument used for analysis was a Waters Acquity system [detectors: Acquity SQD, Acquity ELSD, Acquity PDA] with a Waters ACQUITY UPLC BEH C18 1.7 µm Column. The analysis was performed at a flow rate of 1 mL/min with a linear gradient over 2 min (3 to 99% acetonitrile in water, 0.1% formic acid as modifier). Proton ($^1$H) NMR spectra were recorded on a Bruker Avance 400 spectrometer (400 MHz) using DMSO-$d_6$ or $CDCl_3$ as solvents. Chemical shifts are given in parts per million (ppm) (δ relative to residual solvent peak for $^1$H).

**Compound Synthesis and Characterization**

Epacadostat was synthesized by a modified literature procedure[1]. Crude product was recrystallized from 2:1 water: methanol (v/v) and dried in a 50ºC convection oven to a constant weight to furnish white solid (purity >99%). LCMS $(M + H)^+$: m/z = 438.0, 440.0. $^1$H NMR (400MHz, DMSO-$d_6$): δ 11.51 (s, 1H), 8.90 (s, 1H), 7.17 (dd, J = 8.8Hz, 8.8 Hz, 1H), 7.11 (dd, J = 6.1, 2.7Hz, 1H), 6.76 (m, 1H), 6.71 (t, J = 6.0Hz, 1H), 6.59 (s, 2H), 6.23 (t, J = 6.1Hz, 1H), 3.35 (dt, J = 10.9, 7.0Hz, 2H), 3.10 (dt, J = 12.1, 6.2Hz, 2H).

**(*R*)-3-(4-(Diisobutylamino)-3-((3-(trifluoromethyl)-1,2,4-thiadiazol-5-yl)amino) phenyl) butanoic acid (GSK5628).**

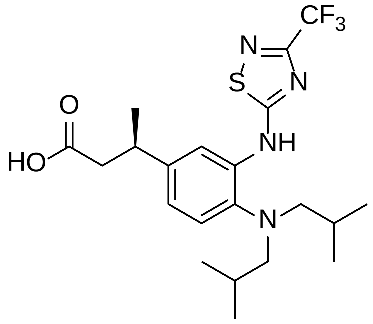


The synthesis of GSK5628 is described in the literature[2].
LCMS $(M + H)^+$: m/z = 459 (ES+)
$^1$H NMR (400 MHz, DMSO-$d_6$) δ ppm 7.57 (br. s., 1 H) 7.16 (d, *J*=8.2 Hz, 1 H) 7.06 (d, *J*=8.1 Hz, 1 H) 3.00 - 3.22 (m, 1 H) 2.72 (d, *J*=6.8 Hz, 4 H) 2.29 - 2.48 (m, 2 H) 1.52 - 1.79 (m, 2 H) 1.19 (d, *J*=6.6 Hz, 3 H) 0.78 (d, *J*=6.4 Hz, 12 H).
$^{13}$C NMR (Chloroform-*d*, 126MHz): δ (ppm) 180.6, 177.1, 159.6, 143.4, 139.4, 135.2, 123.4, 122.4, 118.1, 114.3, 63.9, 42.3, 35.9, 26.2, 21.7, 20.9.

**(R)-3-(4-((5-aminopentyl)(isobutyl)amino)-3-((3-(trifluoromethyl)-1,2,4-thiadiazol-5-yl)amino)phenyl)butanoic acid (GSK7608).**

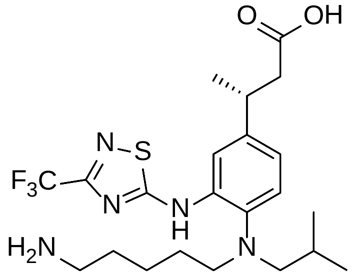


GSK7608 was synthesized according to the procedure described in the literature using the corresponding secondary amine[2].
$^1$H NMR (400 MHz, Chloroform-*d*) δ 7.66 (s, 1H), 7.25 (d, *J*=8.2 Hz, 1H), 7.11 (dd, *J*=8.2 Hz, 1H), 3.29-3.23 (m, 1H), 2.95 (t, *J*=7.3 Hz, 2H), 2.76 (d, *J*=7.0 Hz, 2H), 2.69-2.57 (m, 2H), 2.48 (d, *J*=7.7 Hz, 2H), 1.66 (dt, *J*=13.5, 6.7 Hz, 1H), 1.54-1.35 (m, 4H), 1.35-1.24 (m, 5H), 0.90 (d, *J*=6.6 Hz, 6H).
LC MS *m/z* 488.58 $[M+H]^+$

**(3R)-3-(4-((5-((((E)-cyclooct-4-en-1-yloxy)carbonyl)amino)pentyl)(isobutyl)amino)-3-((3-(trifluoromethyl)-1,2,4-thiadiazol-5-yl)amino)phenyl)butanoic acid (GSK5112)**

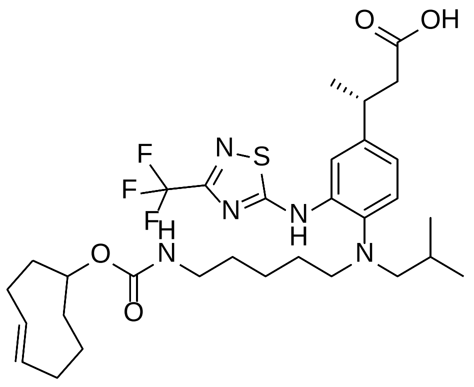

(*R*)-3-(4-((5-amino)pentyl)(isobutyl)amino)-3-((3-(trifluoromethyl)-1,2,5-thiadiazol-5-yl)amino)phenyl)butanoic acid (GSK7608) (8 mg, 0.016 mmol) was dissolved in dimethyl sulfoxide (0.5 mL) and to this was added (*E*)-cyclooct-4-en-1-yl (4-nitrophenyl) carbonate (4.78 mg, 0.016 mmol) and diisopropylethylamine (5.73 µl, 0.033 mmol) . The reaction mixture was stirred for 2 h at room temperature and subsequently purified by preparative HPLC. This afforded (3*R*)-3-(4-((5-((((E)-cyclooct-4-en-1-yloxy)carbonyl)amino)pentyl)(isobutyl)amino)-3-((3-(trifluoromethyl)-1,2,4-thiadiazol-5-yl)amino)phenyl)butanoic acid as a white solid (4 mg, 6.25 µmol, 38.1 % yield). $^1$H NMR (400 MHz, Chloroform-*d*) δ 7.16-7.12 (m, 2H), 6.97 (s, 1H), 6.1 (d, *J* = 20, 1H) 5.49 – 5.35 (m, 2H), 4.82-4.78 (m, 1H), 4.69-4.66 (m, 1H), 3.26-3.24 (m, 1H), 3.10-2.98 (m, 1H), 2.97 – 2.80 (m, 3H), 2.77-2.68 (m, 2H), 2.58 (d, *J* = 8.9 Hz, 3H), 2.41 (q, *J* = 14.1, 13.5 Hz, 1H), 2.27 – 2.09 (m, 4H), 2.05-2.97 (m, 1H), 1.84 – 1.49 (m, 4H), 1.37 – 1.25 (m, 7H), 1.20-1.16 (m, 1H), 1.12 (q, *J* = 14.5, 13.5 Hz, 2H), 0.83 (d, *J* = 6.7 Hz, 6H).
LC MS *m/z* 640.5 $[M+H]^+$ (Rt = 3.76 min).

**(R)-4 & 5-((5-((4-(1-carboxypropan-2-yl)-2-((3-(trifluoromethyl)-1,2,4-thiadiazol-5-yl)amino)phenyl)(isobutyl)amino)pentyl)carbamoyl)-2-(6-hydroxy-3-oxo-3H-xanthen-9-yl)benzoic acids (GSK1051)**

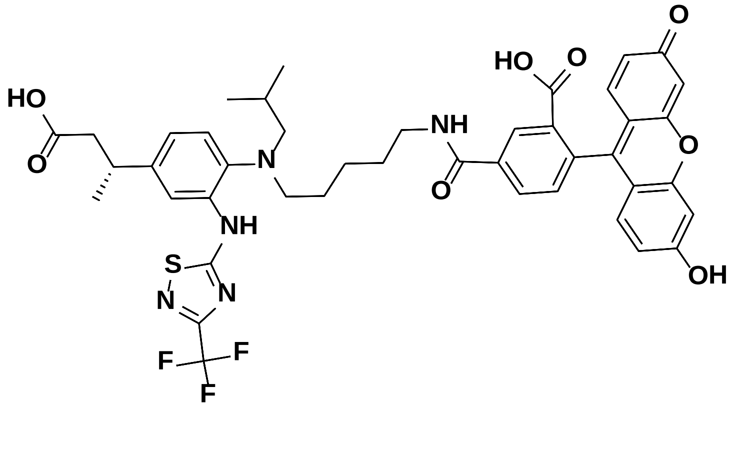

A mixture of (R)-3-(4-((5-aminopentyl)(isobutyl)amino)-3-((3-(trifluoromethyl)-1,2,4-thiadiazol-5-yl)amino)phenyl)butanoic acid (GSK7608) (3.8 mg, 7.79 µmol) 5(6)-Carboxyfluorescein N-hydroxysuccinimide ester (5.8 mg, 0.012 mmol) (mixture of isomers, purchased from Aldrich) in Dichloromethane (DCM) (3 mL) was stirred at room temperature overnight. The reaction mixture was concentrated. The residue was dissolved in MeOH and purified by ISCO 4.3g C18 reverse phase column (mobile phase A: water with 0.1% formic acid; mobile phase B: MeOH; eluted by gradient 40-90% B in

A for 20 column volumes and 90% B in A for 10 column volumes; collected during 9th-12th column volumes) to afford the title compound as a yellow solid.

LCMS: two unequal (~1:2) peaks both having $m/z$ = 846.6 $(M+1)^+$; $t_R$ = 1.27 & 1.29 min; a mixture of (R)-4-((5-((4-(1-carboxypropan-2-yl)-2-((3-(trifluoromethyl)-1,2,4-thiadiazol-5-yl)amino)phenyl)(isobutyl)amino)pentyl)carbamoyl)-2-(6-hydroxy-3-oxo-3H-xanthen-9-yl)benzoic acid and (R)-5-((5-((4-(1-carboxypropan-2-yl)-2-((3-(trifluoromethyl)-1,2,4-thiadiazol-5-yl)amino)phenyl)(isobutyl)amino)pentyl)carbamoyl)-2-(6-hydroxy-3-oxo-3H-xanthen-9-yl)benzoic acid.

$^1$H NMR (400 MHz, Methanol-$d_4$) δ 8.34-8.51 (m, 1H), 8.02-8.25 (m, 1H), 7.55-7.85 (m, 2H), 7.16-7.34 (m, 2H), 7.01-7.18 (m, 1H), 6.45-6.75 (m, 5H), 3.11-3.43 (m, 4H), 2.48-3.00 (m, 5H), 1.19-1.69 (m, 10H), 0.68-1.01 (m, 6H).

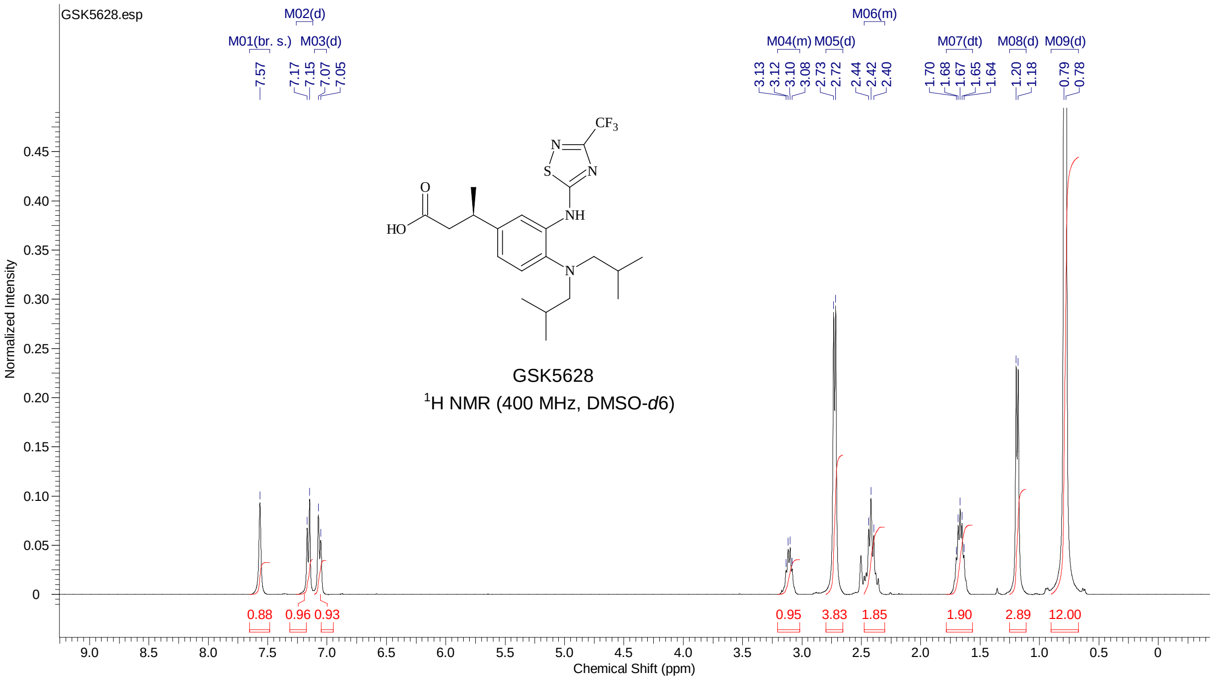

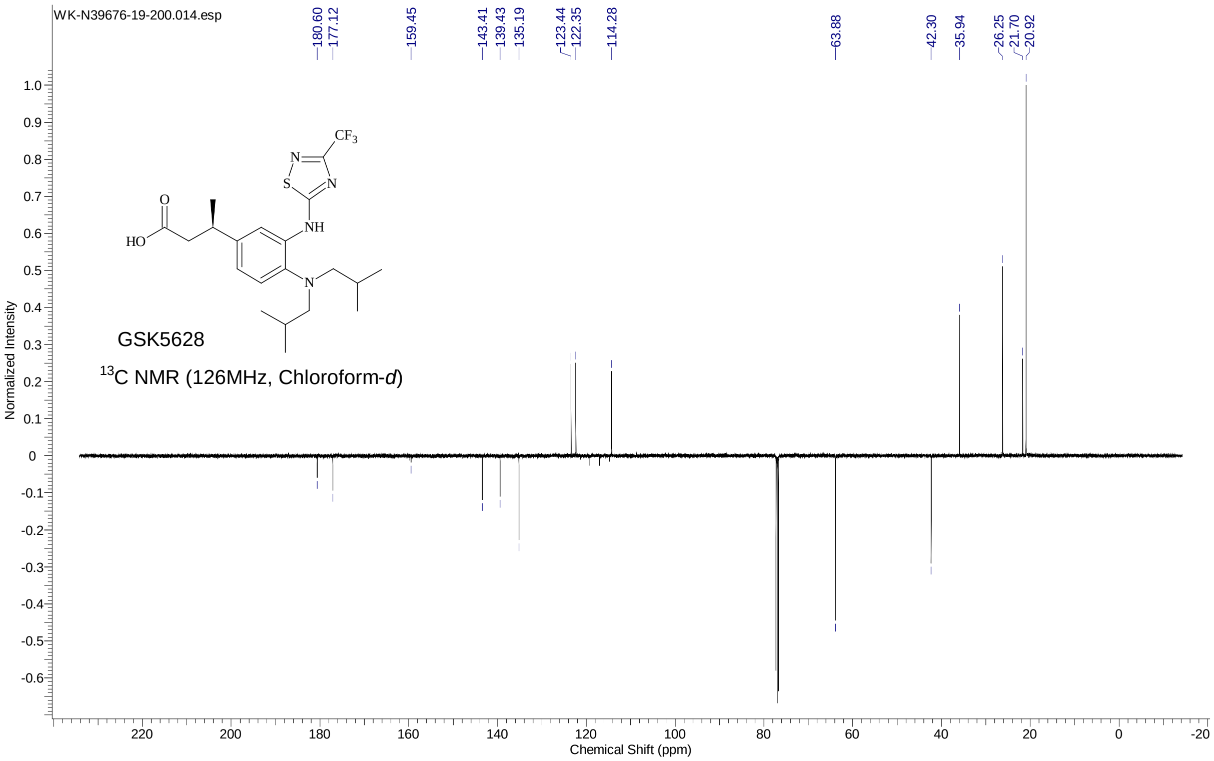


### IDO1 Purification Constructs / Growth Conditions

6H/FLAG/Avi/TEV/IDO1 was assembled from a set of 46 overlapping oligonucleotides and ligated into pET24a. pET24-6H/FLAG/Avi/TEV/IDO1 was freshly transformed into BL21star(DE3), cell growth was carried out at 37°C in *E.coli* production broth (e.g., LB Broth or richer media TB, 2xYT) with carbon source (1% glucose/glycerol), and induced for 20 h at 15°C with 0.5mM IPTG in broth with 1% glucose, 50 µg/ml kanamycin, 100 µM Hemin (Sigma-Aldrich, Cat#. 51280), and 100 µM $FeCl_3$.

### Purification of holo-IDO1

E. coli cell paste expressing 6H/FLAG/Avi/TEV/IDO1 was suspended in 50 mM HEPES, 25 mM imidazole, 500 mM NaCl, 50 mg/L AEBSF (+5 mL protector solution; Sigma-Aldrich, Cat#. 11873601001), pH 8 and lysed with two passes through a Rannie Homogenizer at 10,000 psi. The 0.22µm filtered cell lysate supernatant was loaded onto a Ni Sepharose FF column, resin was washed with 25 mM HEPES, 50 mM imidazole 500 mM NaCl, pH 7.5, and 6H/FLAG/Avi/TEV/IDO1 was eluted with a multi-step gradient to 500 mM imidazole. The Ni elution pool was digested with Tobacco etch virus protease (TEV) and diafiltered prior to flowing through Ni Sepharose FF in the presence of 25 mM imidazole. IDO1 recovered in the Ni unbound pool was diafiltered into 25 mM Tris-Cl, pH 8, loaded onto Q-Sepharose FF (GE Healthcare), and eluted by a 3 column volume linear salt gradient to 0.3M NaCl. IDO1 pools were made based on SDS-PAGE and 405nm/280nm chromatographic profile. The Q Sepharose pool was concentrated and sized on Superdex 200 in 20mM Tris 0.2M NaCl pH=7.4. The monomer peak was pooled. All purification steps performed at 4°C unless otherwise indicated.

**Purification of apo-IDO1**

E.coli cell paste expressing 6H/FLAG/Avi/TEV/IDO1 was suspended in 25 mM Tris, 500 mM NaCl, pH 7.4 10 mM $MgCl_2$[3] and lysed by two passes through a Microfluidizer at 12,000xg. 6H/FLAG/Avi/TEV/IDO1 was batch captured from cell lysate supernatant onto Ni Sepharose Fast Flow (GE Healthcare) and extensively washed with 25 mM Tris, 500 mM NaCl, pH 7.4, with out and then with 30mM imidazole. IDO1 was released from the resin by overnight on column Tobacco etch virus protease cleavage at 4°C. The Ni unbound was dialyzed against 25 mM HEPES 0.25 M NaCl 0.1 mM TCEP. Heme was released from the IDO1 by overnight room temperature incubation with 166 mM Na 2-mercaptoethanesulfonate (MESNA). MESNA forms a temporary thioester with IDO1, impacting heme binding. The apo-IDO1 was concentrated and diafiltered against 25mM Tris pH 8, 0.1 mM TCEP at room temperature prior to flowing through a 5 ml QHP HiTrap (GE Healthcare). The apo-IDO1 in the Q flow through was concentrated and separated by size on Superdex 200 in 20 mM Tris, 0.2 M NaCl pH7.4. The monomer peak was pooled and heme occupancy was determined by comparing the absorbance ratio of 405 nm/280 nm. All purification steps were performed at 4°C unless otherwise indicated.

**Recombinant IDO1 Activity Assay**

Recombinant IDO1 activity was measured largely as previously described[1]. Briefly, 100 nM holo-IDO1 in 100 mM potassium phosphate buffer (pH 7.2) plus 1 mM CHAPS supplemented with 0.5% (v:v) catalase (Sigma 02071) was added (25 µL) to inhibitors in DMSO carrier (0.5 µL DMSO volume) and allowed to incubate for the indicated time and temperature (varied). After the incubation, 4 mM D-TRP, 20 mM L-ascorbic acid and 1 µM methylene blue in 100 mM Potassium Phosphate buffer (pH7.2) plus 1 mM CHAPS were added to the reaction (25 µL). The absorbance at 320 nm was measured at indicated times (typically 0 and 60 minutes) on a Spectramax plate reader. From those measurements, % inhibition was calculated and, where applicable, $IC_{50}$ values were generated (Prism GraphPad). Where indicated, incubation time and temperature, order of addition and reagent composition were varied from this protocol. For **Figure 4B** 50 nM holo-IDO1 and 5 µM GSK5628 were incubated at either 25°C (purple line) or 37°C (orange line) for the indicated times. After the preincubation, 2 mM D-TRP was added to the mixture and the absorbance at 320 nm was read on a Spectramax instrument at 0 and 10 minutes. % inhibition was calculated relative to a similarly incubated DMSO-only control to account for any enzyme degradation. Results are average and standard deviation of N=3 experiments. A single exponential fit was used to generate rate constants at both temperatures, which approximate the limiting apparent heme dissociation rates.

For the reconstitution of apo-IDO1 activity (**Figure 4C**), 100 nM apo-IDO1 in 100 mM potassium phosphate buffer (pH 7.2) was added (25 µL) to varying heme (hemin, Sigma-Aldrich, Cat#. 51280) concentrations in a DMSO carrier (0.5 µL DMSO volume) and allowed to incubate at the indicated temperature for 10 minutes. After the incubation, 4 mM D-TRP, 20 mM L-ascorbic acid and 1 µM Methylene Blue in 100 mM Potassium Phosphate buffer (pH7.2) supplemented with 0.5% (v:v) catalase (Sigma 02071) were added to the reaction (25 µL). The absorbance at 320 nm was measured at 0 and 60 minutes on a spectramax plate reader. 100 nM holo-IDO1 was assayed as a control for 100% reconstituted IDO1 activity. To estimate the affinity of heme for apo-IDO1, results were converted to Abs/min and analyzed using a tight binding fit equation (v = Vmax*((E + S + Km)-sqrt((E + S + Km)^2 - (4*E*S)))/(2*E))[4] with apo-IDO1 (E) constrained at 50 nM, S = heme, Km = heme Kd, v and Vmax expressed as % reconstituted activity.

**Affinity Selection / Mass Spectrometry (AS/MS)**

**Preparation of Resin for Size-Exclusion Chromatography.** 45 g of resin powder (Bio-Rad Bio-Gel® P10 fine resin) were added to a solution of 10 mM Tris, pH 7.5 and 0.02% of sodium azide and slowly mixed until a suspension was formed. The suspension was allowed to swell overnight, and the volume adjusted the day after to 500 mL. On the day of the experiment, 130 µL of Bio-Gel® P10 resin suspension were added to each well of a low protein binding Millipore Multiscreen HTS 384 well HV filter plate (referred here as "size-exclusion plate") with a 0.45 µm Durapore (PVDF) membrane (#MZHCN0W10). The plate was centrifuged at 1000 X g for 2 minutes, and the eluent flow through discarded. 50 µL of 100 mM potassium phosphate (pH 7.2) plus 1 mM CHAPS was then added to the size exclusion plate, which was then centrifuged at 1000 X g for 2 minutes. The wash steps were repeated three additional times to allow resin equilibration. At the end of the washes, 15 uL of 5 µM holo-IDO1 and 100 µM GSK5628 incubated for the indicated time and temperature in 100 mM potassium phosphate (pH 7.2) plus 1 mM CHAPS was transferred to the size-exclusion plate. The size-exclusion plates were centrifuged at 1000 X g for 2 minutes and the flow-through fraction collected into a 384-well clear polypropylene plate containing 5 µL of water. Plates were sealed and stored at -80°C overnight. Finally, plates were centrifuged at 2000 X g for 30 minutes to precipitate protein prior LC-MS analysis. Samples with 100 µM GSK5628 were used as a negative control and no GSK5628 was detected without the presence of holo-IDO1 to allow the compounds passage through the size-exclusion resin.

**Liquid Chromatography-Mass Spectrometry (LC-MS).** All analyses were performed on a Waters Xevo G2S time-of-flight MS interfaced with a Waters Acquity I-Class UPLC and MassLynx 4.1 acquisition software. 10 µL sample injections were made in partial loop-fill mode on a Waters SM-FL autosampler. LC separation used a Thermo Hypersil Gold 20 mm x 2.1 mm column with 1.9 µm particles column, heated to 55° C. The mobile phase consisted of water (solvent A) and acetonitrile (solvent B), each containing 0.1 % formic acid. The LC method consisted of a linear gradient at 1.4 mL\minute from 1-99 % B over 1.8 minutes, a hold for 0.1 minutes, and back to 1 % B in 0.01 minutes. Autosampler time between injections substituted for column re-equilibration. Approximately one-third of the 1.4 mL/minute flow was directed into the mass spectrometer, and the first 0.1 minutes of the gradient containing salts was diverted away. The MS source was operated in positive-ion, centroided acquisition mode with a source temperature of 150° C, capillary voltage 1 kV, cone voltage 80 V, desolvation temperature of 500° C, cone gas 175 L/hr and desolvation gas 1000L/hr. A leucine/enkephalin lock-mass solution was infused to apply automated mass correction to all spectra acquired from 115 to 1300 m/z in 0.1 s. Data processing was accomplished with custom, in-house software. The mass spectrometric, smoothed, extracted ion-chromatogram (XIC) peak areas and heights and retention time for each compound were recorded. Peak area corresponding to GSK5628 are reported and are the mean of N=3 plus SEM.

**Heme Detection**

Free heme in solution was detected with a commercially available hemin detection kit (SIGMA, Cat#. MAK036). 1 µM holo-IDO1 in 100 mM potassium phosphate buffer (pH 7.2) plus 1 mM CHAPS was incubated with 20 µM GSK5628, epacadostat or equivalent volume of DMSO for 120 min at 37°C. The sample was diluted 1000-fold in the provided Hemin Assay Buffer and the Kit reagents added according to the manufacturer's specification. Absorbance at 570 nm was measured on a Spectramax plate reader and these values are reported normalized to a no DMSO protein control. Values are mean of N=3 plus SEM.

### Cellular Thermal Shift Assay

**Constructs and Virus Generation** – Full-length IDO1 sequence consistent with the reference sequence NM_002164/GeneID #3620 with a 10 amino acid spacer and ePL fused to its carboxy terminal was cloned into a modified version of the pHTBV1 BacMam expression vector.[5] Recombinant BacMan viruses were generated as described in the Bac-to-Bac kit instructions (Invitrogen, Carlsbad CA).

HeLa S3 cells were grown in cell culture medium DMEM/F12 containing 10% FBS at 37$^{o}$C with 5% $CO_2$. For BacMam transduction, virus was added to the cells at an approximate cell density of $2e^6$/mL. 24 hours post-transduction, culture medium was replaced with low serum DMEM/F12 containing 1% FBS and treated with 10 µM tool compound or DMSO as a control. Cells were agitated for 3.5 hours by shaking on a Thermo-Shaker PST-100HL. Post incubation, cells were transferred to a 96 well PCR plate, and the samples were heated at 36°C-69°C with 3°C increment for 3 min followed by a 1 min cool down to room temperature, using Veriti 96 well Thermal Cycler from Applied Biosystems. InCELL Pulse reagent mixture (DiscoverX) was added to the the equal volume of cells at the prescribed ratio (four parts substrate, one part EA reagent, one part lysis detergent). Enzymatic turnover was conducted for 3 h at room temperature in a dark location. The plates were then read using an EnVision 2104 Multilabel plate reader (PerkinElmer, Waltham, MA).

### UV Spectrum Study

A UV−visible spectra study was performed to evaluate the binding of the compounds. The IDO1 absorbance spectra were measured using a Spectramax GEMINI XPS for wavelengths between 250 to 600 nm in 1 nm increments. Samples of 10 µM IDO1 or other heme-containing enzymes were placed in a 96 well UV compatible plate, with 10 µM compounds or DMSO were added, and the mixture were incubated at 37°C for 2 hours before the absorbance measurement at room temperature.

### HeLa assay to measure compound potency / cytotoxicity

A suspension of 100,000 cells/mL HeLa cells was prepared in DMEM high glucose media (plus 10% FBS, 1X Penicillin-Streptomycin, GIBCO) with 10 nM (final) human interferon-γ (R&D Systems)and dispensed onto 384-well Greiner clear PP plates (Cat #781280) containing 500 nL of varying concentrations of compound in a DMSO carrier. Cells were covered with Aera Adhesive seals and incubated for 48 hr at 37°C in a 5% $CO_2$ incubator. After incubation, 10 µL of media from each well was precipitated in 40 uL acetonitrile plus internal standard, deuterated kynurenine sulfate (Cambridge Isotope Laboratories), and spun at 2000 rpm for 10 minutes. 10 µL from each well of the precipitation plate was added to 90 µL H2O and spun at 2000 rpm for 10 minutes. This plate was sealed and the levels of KYN were measured on a rapidfire-mass spectrometer (Agilent) with the internal standard used as an injection control. From those measurements, % inhibition was calculated and, where applicable, $IC_{50}$ values were generated (GraphPad Prism). To determine toxicity, 10 µL of Cell-TiterGlo was added to the remaining 90 uL of the HeLa cell plate (all wells except col 18). This was allowed to incubate for 30 minutes and then read on a PerkinElmer ViewLux Microplate Imager.

**Fluorescence polarization (FP) competitive binding assay for apo-IDO1.**

A fluorescent polarization based binding assay was developed to quantify potency of novel small molecule inhibitors binding to apo-IDO1, by competition with a fluorescently labeled ligand (GSK5628 analog GSK1051 labeled with fluorescein, **Figure S10A**). Typically, test compounds (GSK5628, heme (hemin, Sigma-Aldrich, Cat# 51280) and epacadostat) were prepared in 100% DMSO and 100 nL was dispensed into 16 individual wells of a 384 well plate to give final concentrations ranging from 0.3125 nM to 10 μM in 2-fold increments. Ligand in 5 μL assay buffer (100 mM potassium phosphate pH 7.2, 0.02% Pluronic F-127 and 0.005% gelatin) was added and the instrument adjusted to 50 mPs using wells with ligand and buffer only (10 uL volume). Following this adjustment, 5 μL of apo-IDO1 prepared in assay buffer was added to the plate. The plate was inserted into the instrument chamber, which was maintained at the appropriate temperature. Samples were read for up to 60 min on the PHERAstar FS instrument (BMG Labtech) capable of measuring fluorescent polarization (excitation 485 nm; emission 520 nm; 520 nm dichroic). The final concentrations of human apo-IDO1 enzyme and fluorescent ligand in 25 $^{o}$C trials were 10 nM and 5 nM (aproximately 0.5 x Kd; see below), respectively. All FP binding $IC_{50}$ data were normalized to the mean of high and low control wells on each plate. Test compound inhibition was expressed as percent inhibition of internal assay controls. The 5 min read time was used for epacadostat and the 20 min read time was used for heme and GSK5628. Compounds were tested with at least n = 2 in the apo-IDO1 FP binding assay and the mean $IC_{50}$ reported. A four parameter curve fit was used (GraFit; Erithacus Software) to determine the potency of the compounds, where Y is % inhibition, x is inhibitor concentration, 'a' is the minimum (locked at zero), 'b' is the Hill slope, 'c' is the $IC_{50}$ and 'd' is the maximum: $Y=(a-d)/(1+(x/c)^b)+d$ (main body **Figures 4E**). Potency is underestimated due to the relatively high apo-IDO1/probe ratio needed for the FP assay format. Conversion of $IC_{50}$ values to Ki values is estimated by matching $IC_{50}$ values calculated at different Ki's from E, L, and ligand Kd inputs using modifcations of the methods described in Jameson and Mocz[6] and Kuzmic et al[7]. Thus, at 25$^{o}$C, both heme and GSK5628 Ki values are <0.5 nM and at 37$^{o}$C, the heme Ki is ~ 4.1 nM and GSK5628 <0.1 nM by this method.

To determine the Kd of the fluorescent ligand GSK1051, a dual titration of ligand (varied from 1.25 to 120 nM in 2-fold increments as indicated) versus apo-IDO1 (varied from 0.061 to 1000 nM in 2-fold increments at each ligand concentration as indicated) was conducted at ambient and 37$^{o}$C temperatures (upward drift of the ligand/probe FP at 37$^{o}$C was reduced significantly by lowering the Pluronic F-127 to 0.005% and the total DMSO to 0.5%). Data were fitted in GraFit (Erithacus Software), **Figure S12B and C,** to a quadratic solution to binding by anisotropy (r) taking quench upon binding into account: r = ((rfree*f)+(rbound*FR*Q))/(f+(Q*FR)) where rfree is the anisotropy of free ligand, rbound is the anisotropy of bound ligand, Q is bound intensity/free intensity; f = Ftot – FR; Ftot is the varied total ligand concentration; FR = (-(-b + sqrt(sqr(b) - 4 * Ftot * a * Rtot)) / 2); Rtot is the varied total apo-IDO1 concentration, a is the fractional activity; and b = Kd + Ftot + (a*Rtot)[6,7].

The association rate constant for the fluorescent ligand was determined by measuring the decreases in the ligand's fluorescence upon binding to apo-IDO1 in a stopped-flow spectrofluorimeter. GSK1051 was titrated (6.25 to 100 nM final in 2-fold increments) at 2x in buffer (100 mM potassium phosphate pH 7.2). 10 nM apo-IDO1 (final) was prepared in buffer at 2x. Inhibitor and enzyme were rapidly mixed using an Applied Photophysics SX20 stopped-flow spectrometer (fluorescence mode, excitation 493 nm, emission > 530 nm, bandpass filter) at ambient temperature (22$^{o}$C). PM voltage was set to approximately 11 volts determined after mixing equal volumes of 200 nM GSK1051 and buffer. After this adjustment, enzyme and GSK1051 were rapidly mixed and 10,000 data time points were collected for up to 400 seconds for each inhibitor concentration, starting with the lowest. Progress curves were averaged (n=4 to 5) and fitted to a single exponential (with a linear component for photobleaching signal decay at 25°C) and offset to determine the $k_{obs}$ at each inhibitor concentration (**Figure S13A**). In contrast to room temperature the fluorescence increased very slightly at 37°C rather than decreased. The $k_{obs}$ were then plotted as a function of inhibitor concentration and increased linearly with increasing inhibitor concentrations suggesting a simple one-step binding mechanism in the concentration range studied. The slope is equal to an on rate constant for GSK1051 of 1.65 $\mu M^{-1}s^{-1}$ or 1.65 x $10^6$ $M^{-1}s^{-1}$ at 25°C (**Figure S13B**). Using a Kd = 10.6 nM, the calculated $k_{off}$ = 0.0175 $s^{-1}$, which is within 2-fold of the measured value below. Due to the small change in the intrinsic fluorescence at 37°C , the probe $k_{on}$ and $k_{off}$ were also measured by KinTek Explorer[8,9] global fits of the FP time courses at 15, 30, 45 and 60 nM probe in duplicate as described in the next section (**Figure S13D**). See the figure legend for additional details for 37°C data.

The dissociation rate constant for the fluorescent ligand was determined by pre-forming the apo-IDO1-ligand complex with 40 nM final apo-IDO1 and 10 nM final GSK1051 at 25°C and 60 nM final apo-IDO1 and 15 nM final GSK1051 at 37°C in 5 uL followed by addition of excess unlabeled GSK5628 (500 nM final) in 5 uL using a PHERAstar FS equipped with an injector (upward drift of the ligand/probe FP at 37°C was reduced significantly by lowering the Pluronic F-127 to 0.005% and the total DMSO to 0.5%). Data were fitted to a single exponential with an offset term $y = A_0.e^{-kt} + \text{offset}$ (**Figure S13C**). The average $k_{off}$ from triplicate experiments at 25°C was 0.00884 $s^{-1}$; $t_{1/2\ off}$ = 78 sec. Using a Kd = 10.6 nM, the calculated $k_{on}$ = 0.834 $\mu M^{-1}s^{-1}$ or 8.34 x $10^5$ $M^{-1}s^{-1}$, which is within 2-fold of the measured value above. See the figure legend for additional details for 37°C data.

**GSK1051, GSK5628 and heme binding rate constant determinations by fluorescence polarization**

A fluorescence polarization (FP) competitive binding assay was used to determine the on and off rate constants for GSK5628 and heme (hemin, Sigma-Aldrich Cat# 51280) at both 25°C and 37°C, and for GSK1051 at 37°C. This assay utilizes a labelled, reversible ligand (GSK1051 see above) that is competitive with GSK5628 and heme. In a typical experiment, 100 nL of titrated inhibitor in DMSO was dispensed into individual wells of a 384-well plate (0, 1.25 to 20 nM final GSK56280; 0, 0.3125 to 40 nM heme in 9 dilutions). Ligand in 5 µL buffer (100 mM potassium phosphate pH 7.2, 0.02% Pluronic F-127 and 0.005% gelatin) was added to a 384 black low volume Greiner plate (upward drift of the ligand/probe FP at 37°C was reduced significantly by lowering the Pluronic F-127 to 0.005% and the total DMSO to 0.5%). Then 5 µL of enzyme was added to the plate to initiate the reaction using the PHERAstar injector and the plate was immediately monitored kinetically for 50 minutes using a PHERAstar plate reader (excitation 485 nm; emission 520 nm; 520 nm dichroic), with the shortest time interval possible between reads to maximize data density. The final concentrations of enzyme and fluorescent ligand were 10 nM and 5 nM at 25°C (**Figure S14A and S15A**) and 30 nM and 15 nM at 37°C (**Figure S14B and S15B**), respectively. To measure the binding of the probe (GSK1051) at 37°C the method was modified to inject ligand into wells containing apo-IDO1 and to slow down the $k_{obs}$ by lowering the final concentration to 7.5 nM ligand injected at 2x, reducing the dead time to 4 sec and increasing the data density to 1 sec read intervals by reading 2 wells at a time containing 2x apo-IDO1 concentration duplicates (15, 30, 45 and 60 nM final) for 15 min (**Figure S13C**). The FP data was converted to the concentration of enzyme-ligand complex (EL) formed by using equation (**1**) and measuring controls for the polarization of free ($P_f$) and bound ($P_b$) ligand, where P = observed polarization and g = g factor [6,7]. The g factor was set to 1.0 in the calculations since the average measured g-factors at both temperatures were close to 1.0 (0.961 at 25$^{o}$C and 1.01 at 37$^{o}$C). The PHERAstar reader is calibrated to 50 mP for the free ligand. The resulting individual binding curves obtained at each inhibitor (or apo-IDO1) concentration were fitted globally using KinTek Explorer to a competitive model which describes both ligand binding and inhibitor binding equilibria (E + L ⬄ EL and E + I ⬄ EI), where 'E' = enzyme, 'L' = ligand, 'EL' = enzyme ligand complex, 'I' = inhibitor, 'EI' = enzyme inhibitor complex, 'k1' = association rate of ligand, 'k2' = dissociation rate of ligand, 'k3' = association rate of inhibitor, and 'k4' = dissociation rate of inhibitor (see below). An additional term EL ⬄ ELS was added in some experiments to account for minor decay of the signal with 'k5' = the rate of decay and 'k6' locked at zero. Total E, L and I were constrained inputs for each experiment at each temperature, and global fits were obtained for the indicated time courses. Multiple iterations of fitting were utilized until rates converged on a global minimum.

$$[EL] = \frac{(3-P_b)(P-P_f)}{(3-P)(P_b-P_f) + (g-1)(3-P_f)(P_b-P)} \qquad (1)$$

$$E + L \underset{k_2}{\overset{k_1}{\rightleftharpoons}} EL$$

$$E + I \underset{k_4}{\overset{k_3}{\rightleftharpoons}} EI$$

**GSK5628 on rate constant and Kd determinations at 22$^{o}$C and 37$^{o}$C using intrinsic apo-IDO1 Trp fluorescence quenching by stopped-flow spectrofluorimetry and equilibrium titrations**

To determine the on rate constant of GSK5628 accurately, decreases in the intrinsic tryptophan fluorescence of apo-IDO1 were measured upon inhibitor binding. GSK5628 was titrated (0.0625 to 4 μM final; epacadostat: 0.5 and 1 μM final at 22$^{o}$C only) at 2x in buffer (100 mM potassium phosphate pH 7.2 with 1 mM CHAPS and 2% DMSO). 50 nM apo-IDO1 (final; 250 nM for **Figure 4D**) was prepared in buffer (no DMSO) at 2x. Inhibitor and enzyme were rapidly mixed 1:1 using an Applied Photophysics SX20 stopped-flow spectrometer (fluorescence mode, excitation 285 nm, emission >320 nm, bandpass filter) equipped with a water bath to maintain temperature at ambient (22$^{o}$C) or 37$^{o}$C. PM voltage was set to approximately 80% of auto PM determined after mixing equal volumes of apo-IDO1 and buffer. After determining this baseline, enzyme and inhibitor were rapidly mixed and 10,000 data time points were collected for up to 20 seconds for each inhibitor concentration, starting with the lowest. Progress curves were averaged (n=3) and fitted to a double exponential to determine the $k_{1obs}$ and $k_{2obs}$ at each inhibitor concentration. Both $k_{obs}$ were then plotted as a function of inhibitor concentration and increased linearly with increasing inhibitor concentrations suggesting simple one-step binding mechanisms in the concentration ranges studied. The slope of the linear fit is equal to the on rate constant for GSK5628 (**Figure S9**).

The Kd for GSK5628 binding to apo-IDO1 at ambient and 37$^{o}$C temperatures was also determined by equilibrium titration of 50 nM apo-IDO1 with varied GSK5628 (14 concentrations from 12 to 500 nM) in 100 mM phosphate buffer, pH 7.2 with 1% DMSO in a 1 mL quartz cuvette. The quenching of the intrinsic trp fluorescence was monitored in a Molecular Devices M2 plate reader with excitation at 285 nm, emission at 365 nm and a cut off at 365 nm (average of 4 reads at each concentration). GSK5628 was diluted in buffer containing 50 nM apo-IDO1 to avoid dilution of the protein and a buffer control at each GSK5628 concentration (average of 4 reads) was subtracted from the protein titrations. Net average fluorescence was fitted to a tight binding equation: A-((A-B)*((K+E+X)-SQRT((K+E+X)^2-4*E*X)))/(2*E)[4], where A is the initial fluorescence intensity, B is the fluorescence of the bound complex, K is the Kd, E is the concentration of apo-IDO1 binding sites, and X is the concentration of GSK5628 (**Figure S10**: (**A**) ambient ~ 22$^{o}$C and (**B**) 37$^{o}$C).

**HDX-MS**

Prior to the experiment, 2 µM IDO1 was incubated with and without 200 µM GSK5628 for 2 hours at 37 °C in 20 mM Tris, pH 7.5, 100 mM NaCl. All following manipulations were conducted with a LEAP PAL HDX in line with an Agilent LC system followed by a Thermo Scientific Orbitrap LTQ-XL mass spectrometer. The chromatography was performed at 4 °C. To initiate the HDX reaction, 2 µL of the prepared sample was mixed with 18 µL unbuffered $D_2O$ and incubated for 30, 100, and 300 s at 23 °C. The reaction was quenched at 0 °C by adding 35 µL of 3.2 M GuHCl, 0.8 % formic acid prior to injection into an in-line protease column (Waters Enzymate BEH pepsin column; 2.1 mm X 30 mm) equilibrated in Buffer A (0.05 % trifluoroacetic acid, 0.1 % formic acid in water). Digest was conducted at 100 µL/min and the resulting peptides were trapped and desalted (Optimize Technologies 3 um EXP C18 Trap column cartridge, 1.0 mm X 5 mm). The peptides were separated and eluted at 10 µL/min using a microbore HPLC column (Agilent ZORBAX SB-C18; 3.5um; 0.5 mm X 35 mm) with a gradient running 10-50 % Buffer B (0.05 % trifluoroacetic acid, 0.1 % formic acid, 5 % water, 95 % acetonitrile) over 14.1 minutes. Peptides were identified via a Mascot search using MS/MS data collected on samples diluted in $H_2O$ instead of $D_2O$. All data analysis was conducted with HDExaminer (Sierra Analytics, Inc). The resulting heat map was plotted onto the epacadostat structure (PBD ID 5WN8) in Pymol.

**Cellular wash-out assays (KYN Level Detection)**

A suspension of 100,000 cells/ml HeLa cells was prepared in DMEM high glucose media, which contains 16 mg/L L-Trp, and 0.1mg/L ferric nitrate (plus 10% FBS, 1X Penicillin-Streptomycin, GIBCO) with 10 nM human interferon-γ (R&D Systems) and 1 µM DMSO, GSK5628 or epacadostat. 1.0 mL was dispensed onto 12-well Greiner clear PP plates and incubated for 24 hr at 37**°**C in a 5% $CO_2$ incubator. After incubation, 10 µL of media from each well was precipitated in 40 µL acetonitrile plus internal standard, Deuterated L-Kynurenine Sulfate (Cambridge Isotope Laboratories), and spun at 2000 rpm for 10 minutes. 10 µL from each well of the precipitation plate was added to 90 µL H2O and spun at at 2000 rpm for 10 minutes. This plate was sealed and the levels of KYN were measured on a rapidfire-mass spec (Agilent) with the internal standard used as an injection control. In order to remove the equilibrated inhibitor population and to prevent new IDO1 synthesis, the remaining media was aspirated and the cells were subsequently washed with DMEM high glucose media (plus 10% FBS, 1X Penicillin-Streptomycin, GIBCO) supplemented with 20 µM Cycloheximide. Wash was repeated 2X times and a 60 minute incubation prior to the final wash was added (See scheme in **Figure S18**). After the final wash, cells were incubated 22 hr at 37**°**C in a 5% $CO_2$ incubator. After incubation, 10 µL of media from each well was checked for KYN levels as described above and cells were subsequently lysed in 1X RIPA buffer (Cell signalling) containing protease inhibitors (Pierce). Cell lysates were analyzed by western blotting to determine protein levels of Actin and IDO1. KYN levels were reported as % IDO1 activity normalized to an untreated (no DMSO) control cell population. Reported values are N=3 and include SEM calculations.

**Cellular competition assays with click probe**

HeLa cells in MEM medium supplemented with 1% puryvat, 1 % NAA and 10% FBS (all from Gibco) were plated on an 8-well Labtek (Thermo Scientific Nunc) tissue culture dishes (per well 200 µl of 0.12 x10^6 cells/ml) and were left overnight in a humidified 37°C, 5% $CO_2$ incubator to adhere. Media was then removed completely by aspiration and replaced with fresh media (200 µl/well) containing 10 nM IFNγ (Sigma) and cells were incubated for another 24 h before treatment with compound. Media was removed by aspiration from the cells and 150 µl fresh media were added into each well. Next, 50 µl of DMSO solution (Sigma) or 50 µl of 4x compound stock solution (GSK5628 or epacadostat, final concentration 5 µM) were added. After 3 hrs of incubation, 20 µl of 10x TCO-probe (GSK5112, final concentration 150 nM) solution were added and cells were then incubated for 1 hr more. After incubation with the compounds media was removed and cells were washed with 200 µl of PBS (Gibco) per well. The cells were then fixed in 4% paraformaldehyde (Electron Microscopy Science) in PBS for 10 min, washed 3 times with PBS and finally permeabilized with 0.5% Triton (Sigma) in PBS. After 3 washes with PBST (0.1% Tween, Sigma, in PBS) cells were incubated with 100 nM Tetrazine-Cy5 (Jena Bioscience) for 5 min at RT. Cells were washed 5 times with PBST and Hoechst staining (Life Technologies) was performed for 10 min at RT. Finally, dishes were washed 2 times with PBST and once with PBS and imaged directly. The image acquisitions were carried out using a Zeiss LSM 780 or Zeiss LSM 780 NLO using 40x or 63x oil objectives. The data was analyzed using ImageJ (1.45s) and CellProfiler (2.1.1) by measuring mean fluorescence intensity in the Cy5 channel (633 nm, corresponding to the signal of click-probe) at single cell level (defined by nuclear staining, 405 nm). Per experiment at least 6 fields of view were imaged corresponding to at least 90 cells being quantified for each condition.

**Cellular wash-out assays with click probe (imaging)**

Concentration of Hela was adjusted to $0.08x10^6$ cells/ml and 200 µl of diluted cells were plated per well on an 8-well Labtek tissue culture dishes. Cells were left for 4 hr at 37°C, 5% $CO_2$ to adhere. Media was removed completely by aspiration and replaced with fresh media (200 µl/well) containing 10 nM IFNγ. Next, 50 µl of DMSO solution or 50 µl of 5x compound stock solution (GSK5628, final 1 µM) was added (all stock solutions were prepared in media + IFNγ) and cells were incubated for 24 h before the wash-out procedure started. Media was removed by aspiration and cells were washed twice, each time with 200 µl of fresh media. Next, 200 µl of media containing cycloheximide (20 uM, Sigma) were added and cells were incubated for 1 hour. Cells were washed once again with 200 µl of media containing cycloheximide and then incubated with 200 µl of media containing cycloheximide for another 22 hr at 37°C, 5% $CO_2$. After this time, 50 µl of DMSO solution or 50 µl of 5x compound stock solution (GSK5628 or epacadostat, final 1 µM) was added (all stock solutions were prepared in media with cycloheximide and cell were incubated for 3hrs. Then, 25 µl of 10x TCO-probe (GSK5112, final 0.15 µM) solution were added and cells were incubated for another 1h at 37°C, 5% $CO_2$. The fixation, staining, imaging and data analysis has been done as described above. Per experiment at least 5 fields of view were imaged corresponding to at least 65 cells being quantified for each condition.

### Supplementary Figures

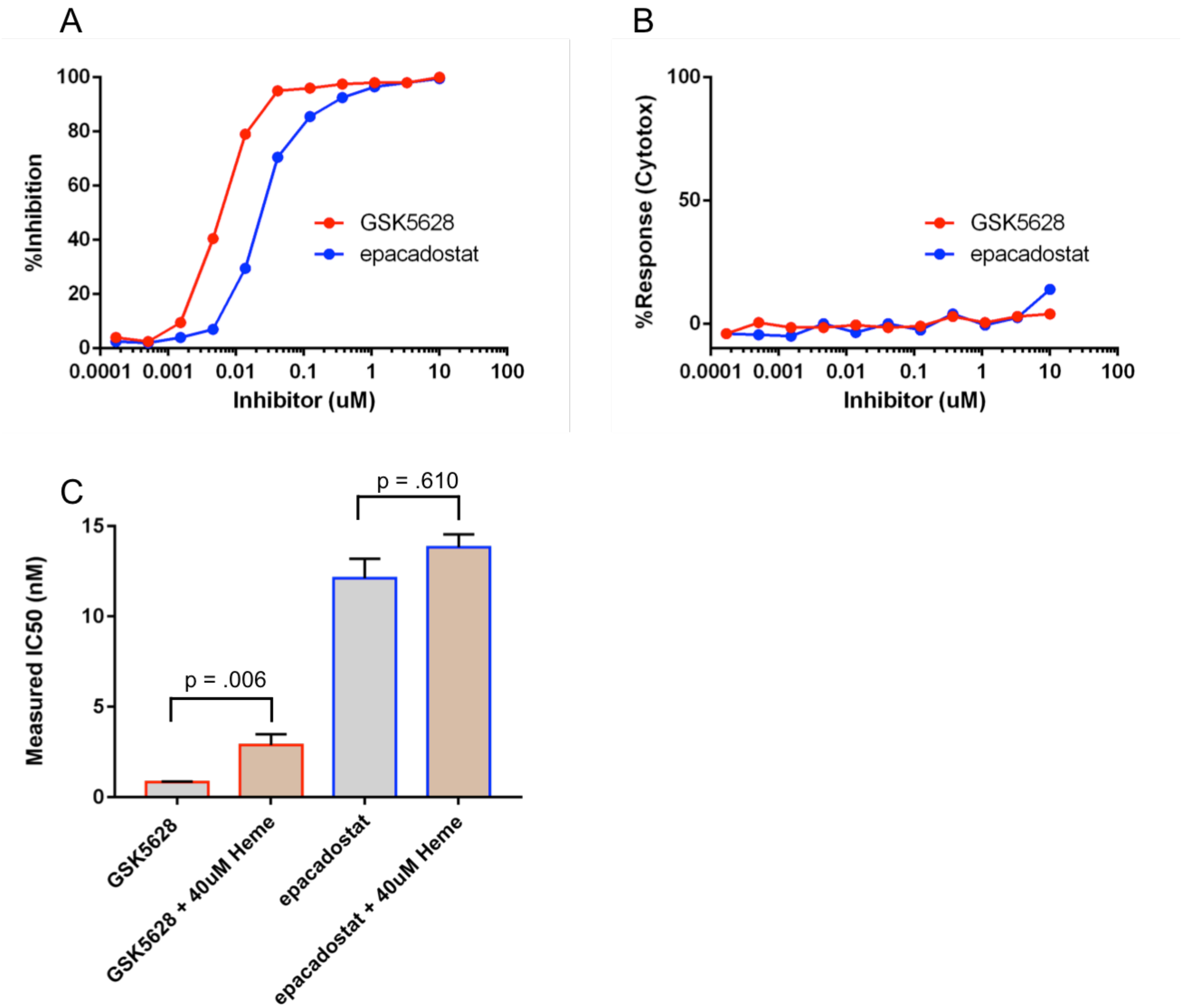


**Figure S1**

Testing the cellular potency and toxicity of GSK5628 and epacadostat. HeLa cells were stimulated with interferon-gamma to stimulate IDO1 production and incubated with varying concentrations (10 µM to 0.17 nM) of GSK5628 (red lines) or epacadostat (blue lines). After a 48 hr incubation, KYN production was measured on a rapidfire mass-spec instrument and transformed into % Inhibition measurements **(A).** These results show that GSK5628 and epacadostat both inhibit IDO1 (epacadostat: 24 nM $IC_{50}$. GSK5628: 5.9 nM $IC_{50}$). Conversely, addition of CellTiter-Glo reagent followed by luminescence measurements shows that neither compound has a significant increase in cytotoxicity at all concentrations tested **(B).** Addition of 40 µM exogenous heme (hemin, Sigma-Aldrich Cat# 51280) to the growth media significantly shifted the measured potency of GSK5628 but not epacadostat **(C)**. Average of 4 independent measurements plus SEM are shown. Statistical significance was calculated using a 2-sample t-test.

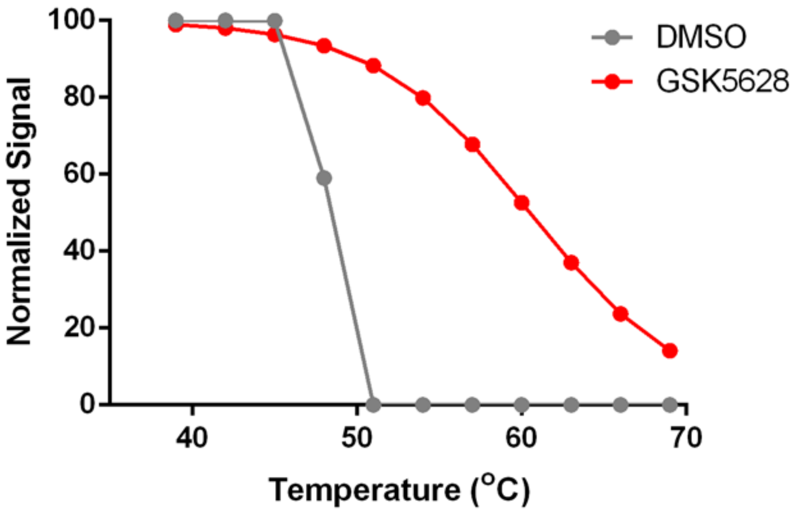


**Figure S2**

Hela cells transduced with an ePL-tagged IDO1 are treated with 10 µM GSK5628 (red line) or DMSO vehicle (grey line), then subjected to different temperatures. IDO1 protein is measured with InCell Hunter detection reagents and levels reported are normalized to cells treated at room temperature.

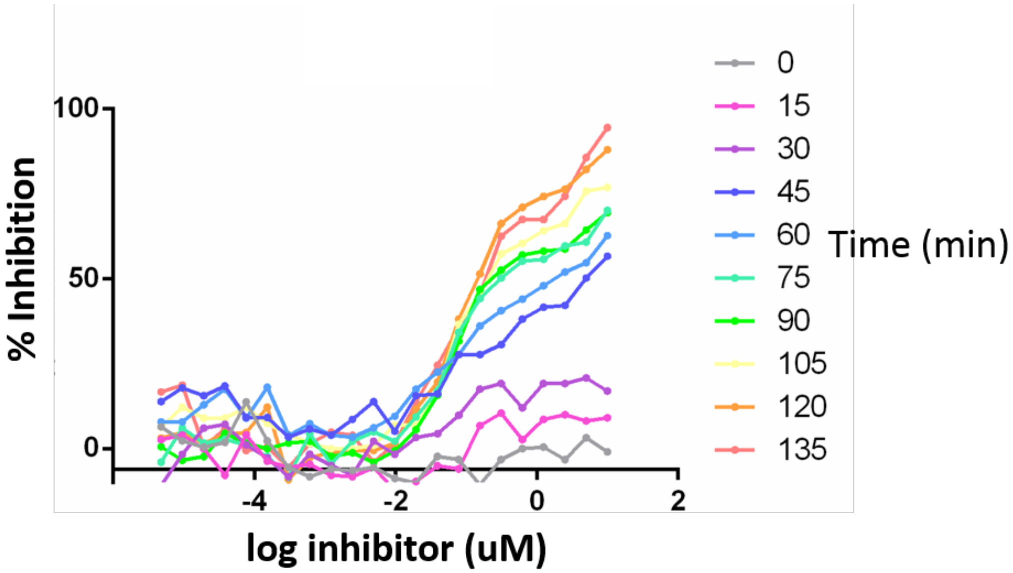


**Figure S3**

GSK5628 shows increased potency with increasing preincubation time. Varying concentrations of GSK5628 were preincubated (before addition of substrate) with 50 nM IDO1 (holo-IDO1) for indicated time at 37°C. After the indicated times, 2 mM D-TRP substrate was added and the absorbance at 320 nm measured at 0 and 60 minutes post substrate addition. From those measurements, % inhibition was calculated and plotted. These results show that, as the preincubation time is increased, the measured potency of 5628A also increases.

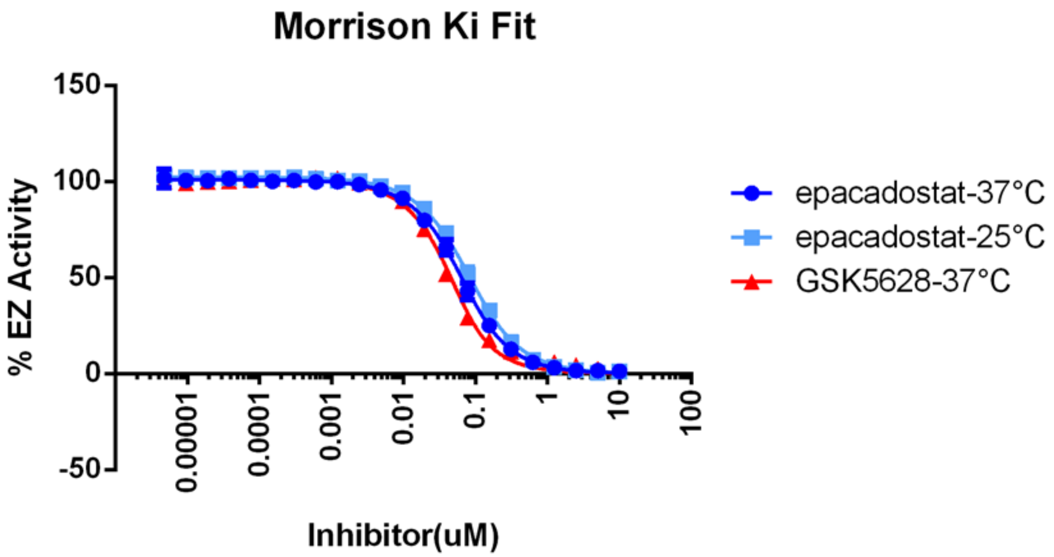


| | epacadostat-37°C | epacadostat-25°C | GSK5628-37°C |
|---|---|---|---|
| Best-fit values | | | |
| Vo | 100.9 | 102.3 | 101.5 |
| Et | = 0.0500 | = 0.0500 | = 0.0500 |
| Ki | 0.01958 | 0.02869 | 0.01014 |
| S | = 2000 | = 2000 | = 2000 |
| Km | = 2100 | = 2100 | = 2100 |

**Figure S4**

Tight binding fit (Morrison Equation[4]) of data from **Figure 2**. Data showing robust inhibition from **Figures 2A and 2B** was converted to %EZ (enzyme) activity and analyzed using the Morrison equation on GraphPad Prism software. Parameters contrained were enzyme catalytic sites at 50 nM, substrate concentration at 2.0 mM and substrate Km at 2.1 mM. Resulting Ki's are reported in micromolar. The Ki values at 37ºC for GSK5628 are within 2-fold of the values obtained by titration of apo-IDO1 TRP fluorescence (**Figure S10**).

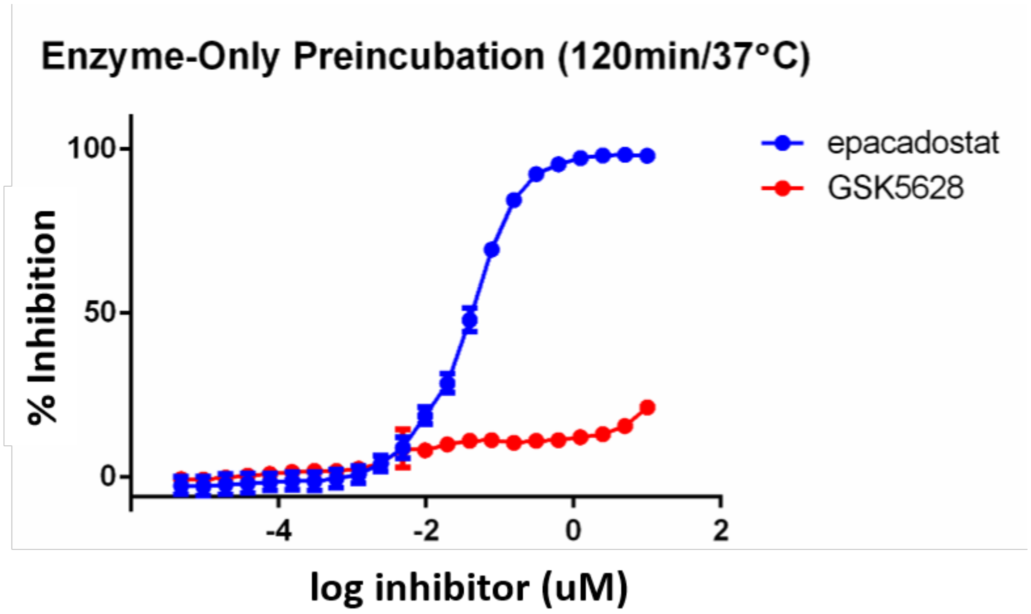


**Figure S5**

Preincubation of IDO1 alone does not cause GSK5628A inhibition. 50 nM holo-IDO1 was incubated for 120 min at 37°C in the absence of any inhibitors. After the incubation, enzyme was added to varying concentrations of GSK5628 or epacadostat. Within 5 minutes, D-TRP substrate was added and the absorbance at 320 nm measured at 0 and 60 minutes post substrate addition. From those measurements, % inhibition was calculated and plotted. These results show that preincubation of IDO1 for 120 min in the absence of GSK5628 does not cause an increase of inhibition when both IDO1 and GSK5628A are preincubated together, as can be seen in **Figure S3**. The inhibition of epacadostat is unaffected. Results are the average and standard deviation of N=3 experiments.

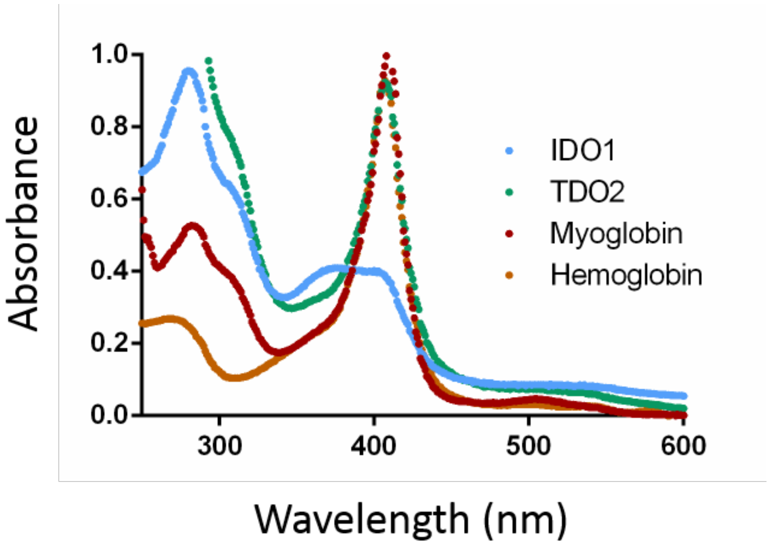


**Figure S6**

Absorbance scans of various heme-containing proteins after GSK5628 incubation show no observed loss of the Soret band except IDO1 (blue). Hemoglobin was obtained from Abcam (Cat# ab77858), myoglobin was obtained from Sigma-Aldrich (Cat# M1882), IDO1 and TDO2 were made in house.

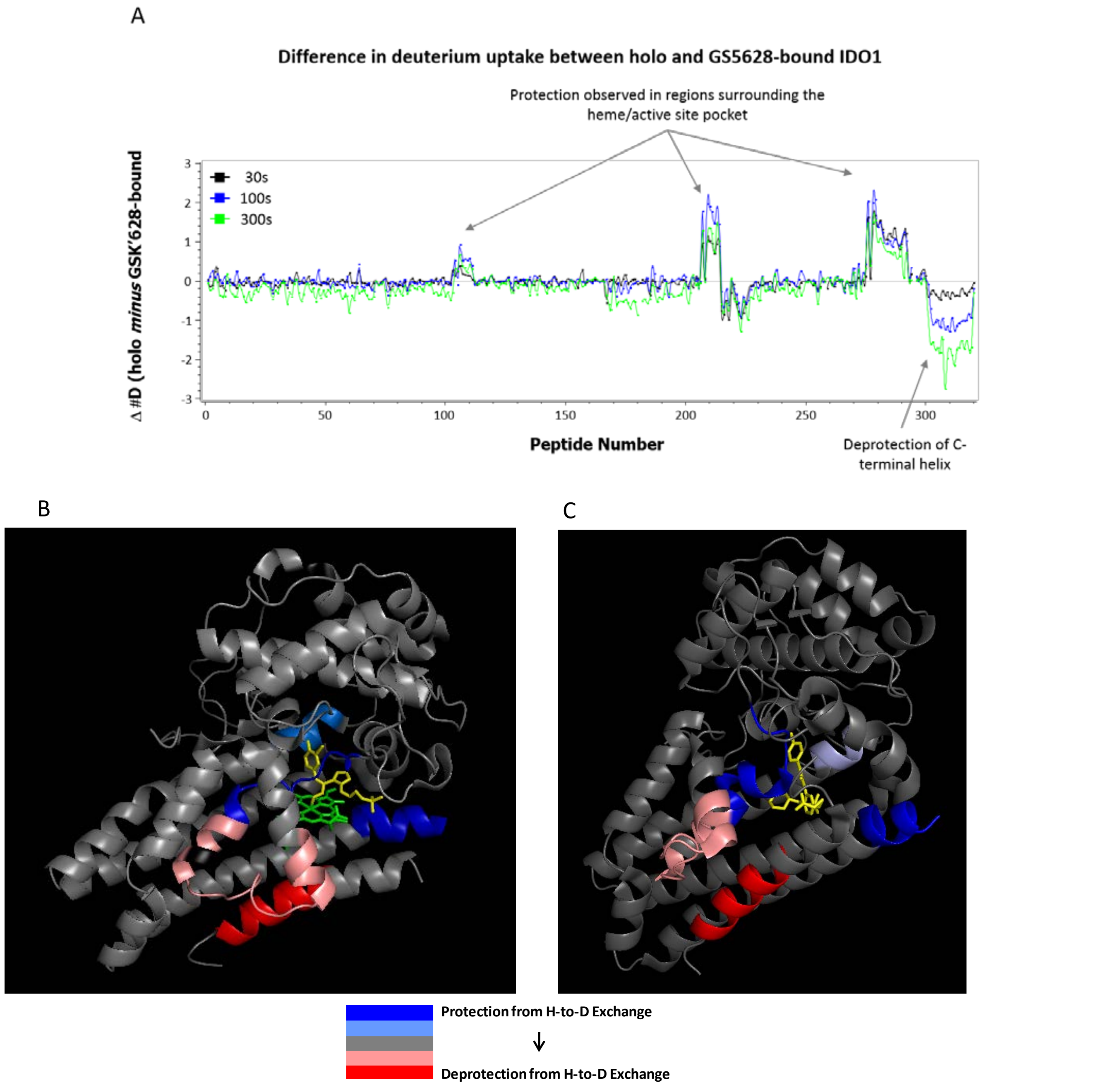


**Figure S7**

HDX-MS studies of holo and GSK5628-bound IDO1. In order to determine the structural effects of GSK5628 on IDO1, HDX-MS studies were conducted on the enzyme after a two hour incubation at 37˚C with and without compound. **(A)** Differential exchange, in deuterons, between holo and compound-bound IDO1 for three deuteration time points (30, 100 and 300 s). The bottom axis is shown as peptide number along the protein sequence. A positive difference indicates that the compound induced protection in a region, whereas a negative difference shows deprotection. **(B and C)** The differential exchange plotted in **(A)** was deconvoluted on a per residue basis and mapped onto the epacadostat-bound IDO1 structure **(B)** (PBD ID 5WN8)[10] and onto the BMS-978587 bound apo-IDO1 structure **(C)**(PDB ID 6AZV)[11]. The heme is colored green **(B),** epacadostat **(B)** and BMS-978587 **(C)** are yellow. Residues that do not show a significant difference in exchange between apo and bound are colored in gray while residues with no exchange information are in black.

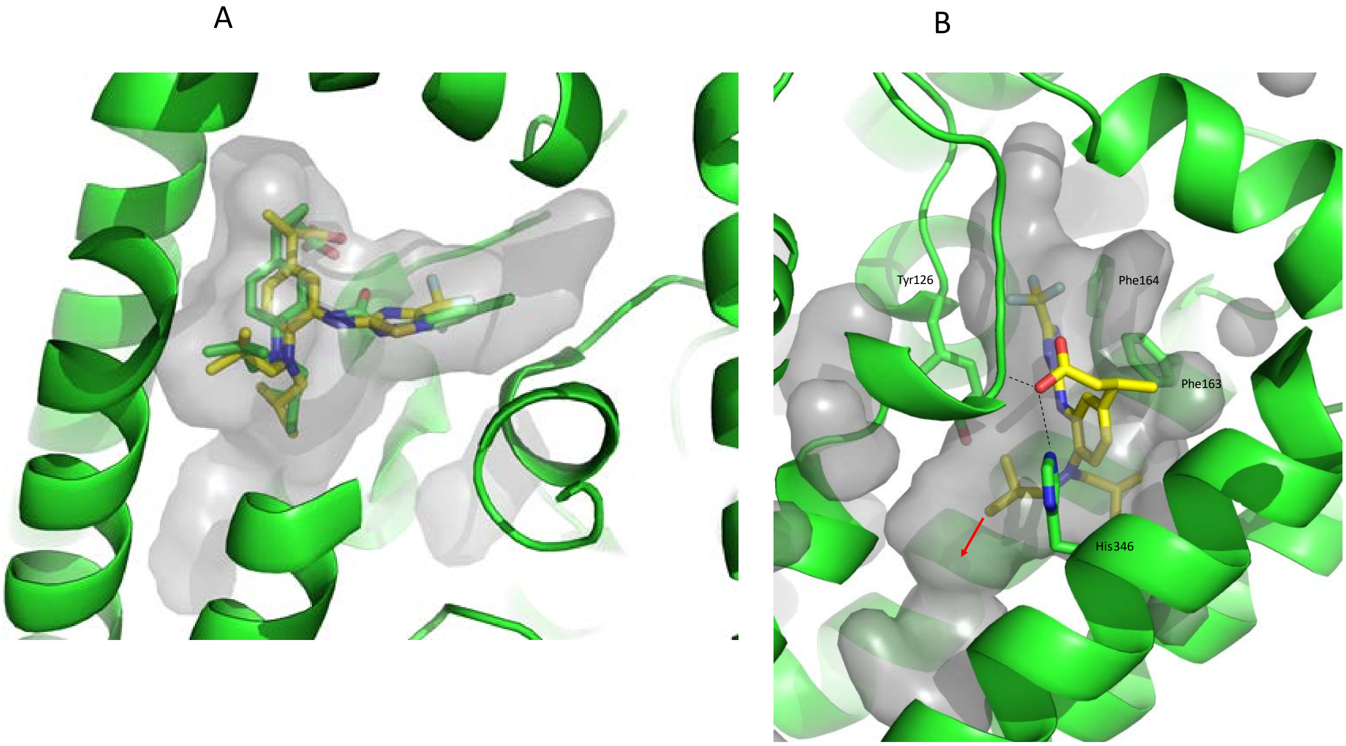


**Figure S8**

Docking models of GSK5628 with apo-IDO1. **(A)** GSK5628 (yellow) docked into the active site of apo-IDO1 (PDB ID 6AZV)**[11]** and overlayed with BMS-978587 (green) showing that GSK5628 could easily be accommodated within the apo binding site. (**B**) GSK5628 docked into the active site of apo-IDO1 (PDB ID 6AZV)**[11]**. The surface of the pocket is highlighted in grey. The GSK5628 carboxylate group is within hydrogen bonding distance of the side chain of His346 and backbone NH of Asp264. A red arrow indicates the position modified in GSK1051 and GSK5112. A tunnel out to solvent exits at this position and could accommodate the linkers and fluorophore of GSK1051 and click probe of GSK5112.

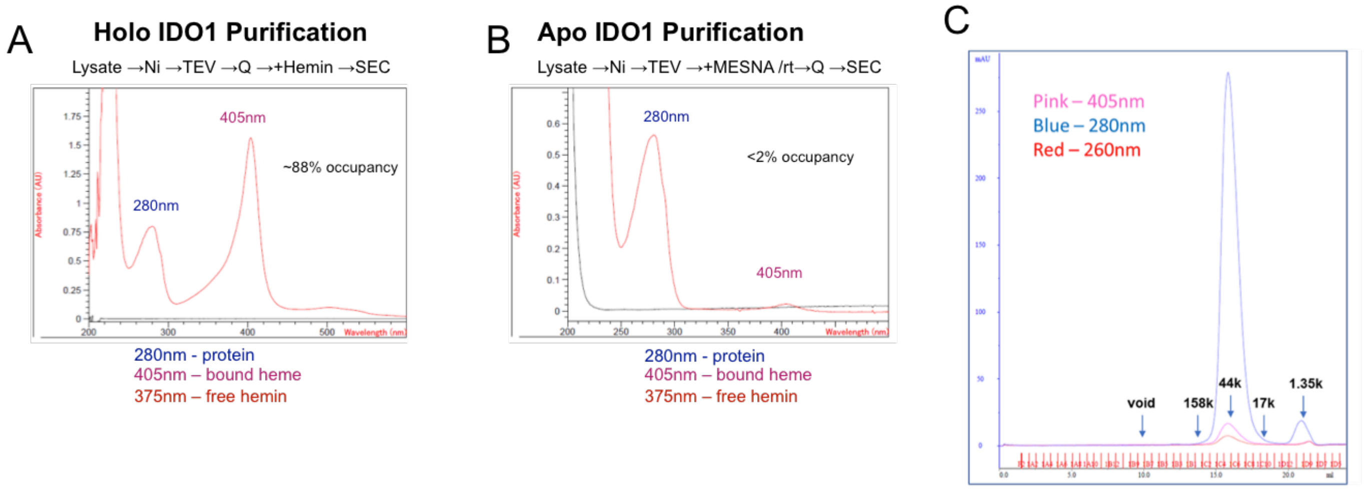


**Figure S9**

Characterization of apo-IDO1. **(A)** The absorbance spectrum of purified holo IDO1 shows that it contains a mixture of heme-bound and heme-free forms of IDO1, with ~88% of the protein heme bound. (**B**) The absorbance spectrum of purified apo-IDO1 shows that it contains a very minimal amount of heme-bound IDO1. **(C)** SEC chromatogram of the apo-IDO1 preparation shows that the apo form of IDO1 does not contain any aggregate peaks. LC-MS analysis of the monomer peak gave the expected m/e ratio for the apo-IDO1 peptide. Based on A405/A28 ratios for apo- and holo-IDO1, apo has ~1.8% residual heme bound. Treatment of the apo-IDO1 pool with 100µM hemin overnight at 4˚C followed by centrifugation and re-scan restored the ratio to ~93% occupancy by heme. No visible precipitate was observed suggesting minimal aggregation of the reconstituted apo-IDO1.

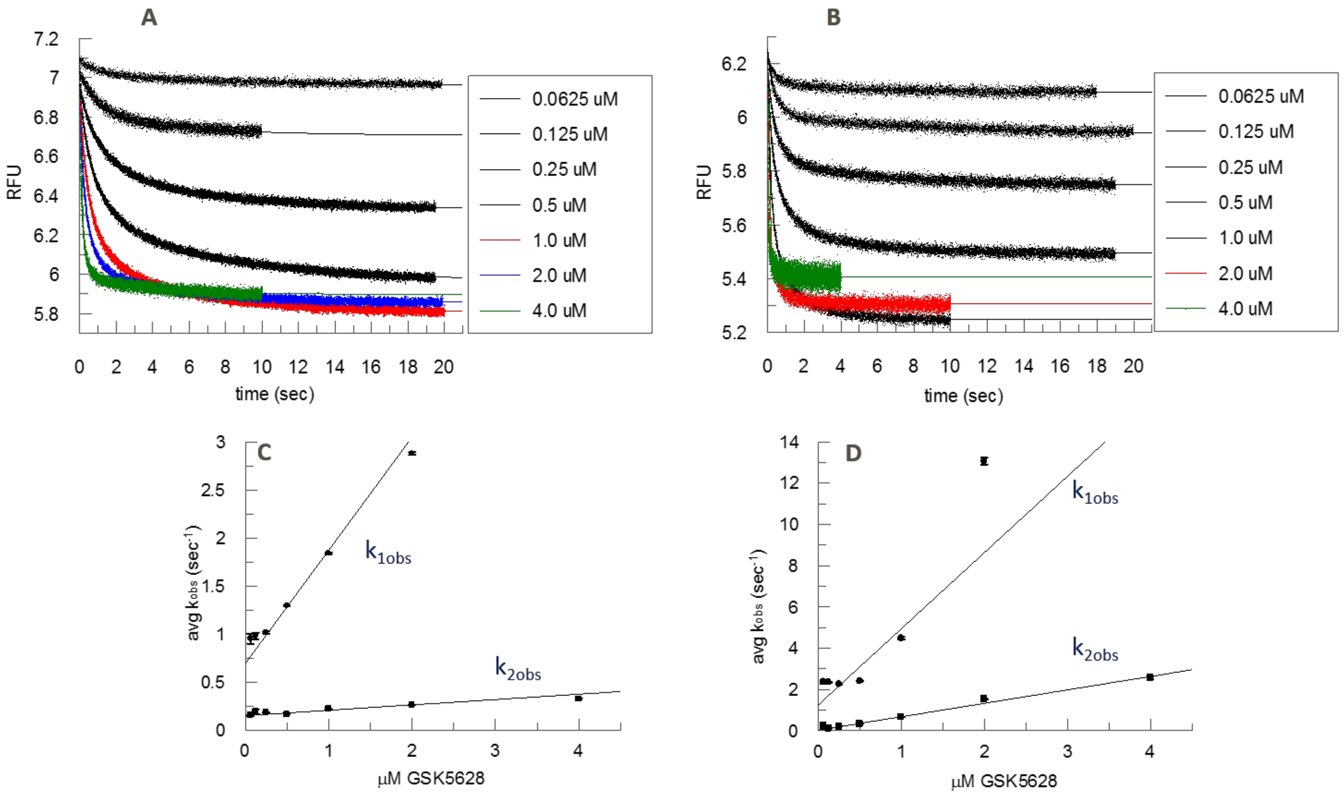


**Figure S10**

Representative stopped-flow time courses (avg n=3; **A** and **B**) of the quenching of intrinsic apo-IDO1 (50 nM) trp fluorescence fit to double exponentials and linear replots (**C** and **D**) of $k_{1obs}$ and $k_{2obs}$ versus GSK5628 concentrations at 22°C (**A** and **C**) and 37°C (**B** and **D**) to determine $k_{on}$ from the replot slopes and demonstrating that direct binding to apo-IDO1 is rapid regardless of temperature. Replots are linear fits of the averages of n=3 runs and the standard deviations using GraFit (Erithacus Software). Values for $k_{on}$ (slopes) are 1.2 ± 0.073 $\mu M^{-1}s^{-1}$ for $k_{1obs}$ (r = 0.990) and 0.056 ± 0.011 $\mu M^{-1}s^{-1}$ for $k_{2obs}$ (r = 0.916) at 22°C and 3.7 ± 1.4 $\mu M^{-1}s^{-1}$ for $k_{1obs}$ (r = 0.773) and 0.65 ± 0.036 $\mu M^{-1}s^{-1}$ for $k_{2obs}$ (r = 0.994) at 37°C. Intercepts poorly define $k_{off}$ due to extreme extrapolation from the lowest concentrations to the Kd's.

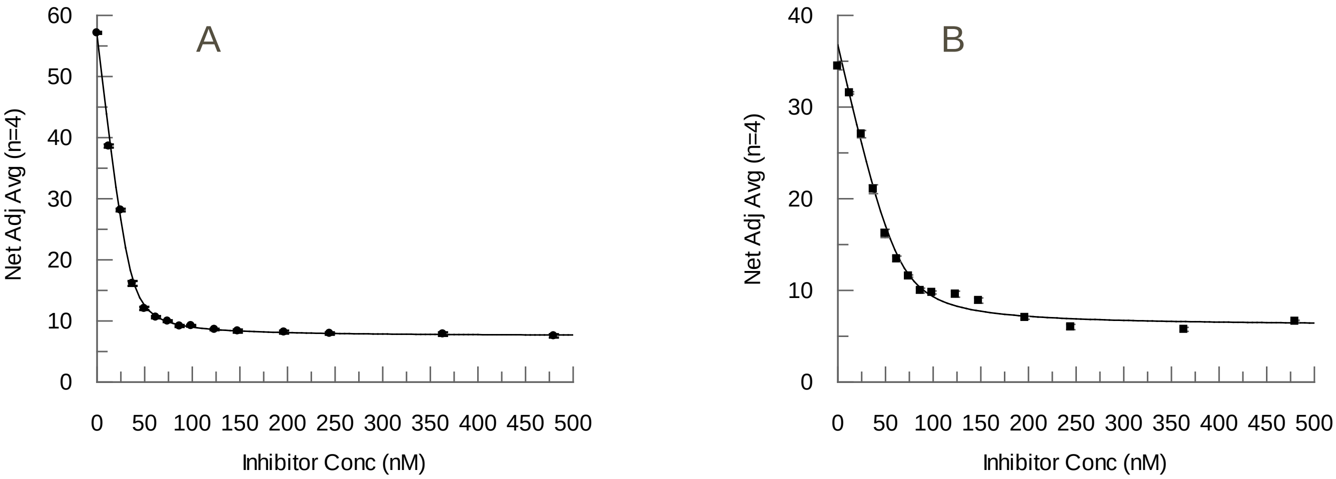


**Figure S11**

Equilibrium titrations of the quenching of intrinsic apo-IDO1 trp fluorescence with GSK5628 at 25$^{o}$C (**A**) and 37$^{o}$C (**B**) to determine an accurate Kd at both temperatures. Fitted parameters shown by the solid lines are (**A**): Kd = 2.1 ± 0.51 nM, E = 35 ± 1.7 nM, A = 57 ± 0.57 RFU and B = 7.5 ± 0.39 RFU; (**B**): Kd = 5.0 ± 1.6 nM, E = 64 ± 0.47 nM, A = 37 ± 0.77 RFU and B = 6.1 ± 0.38 RFU.

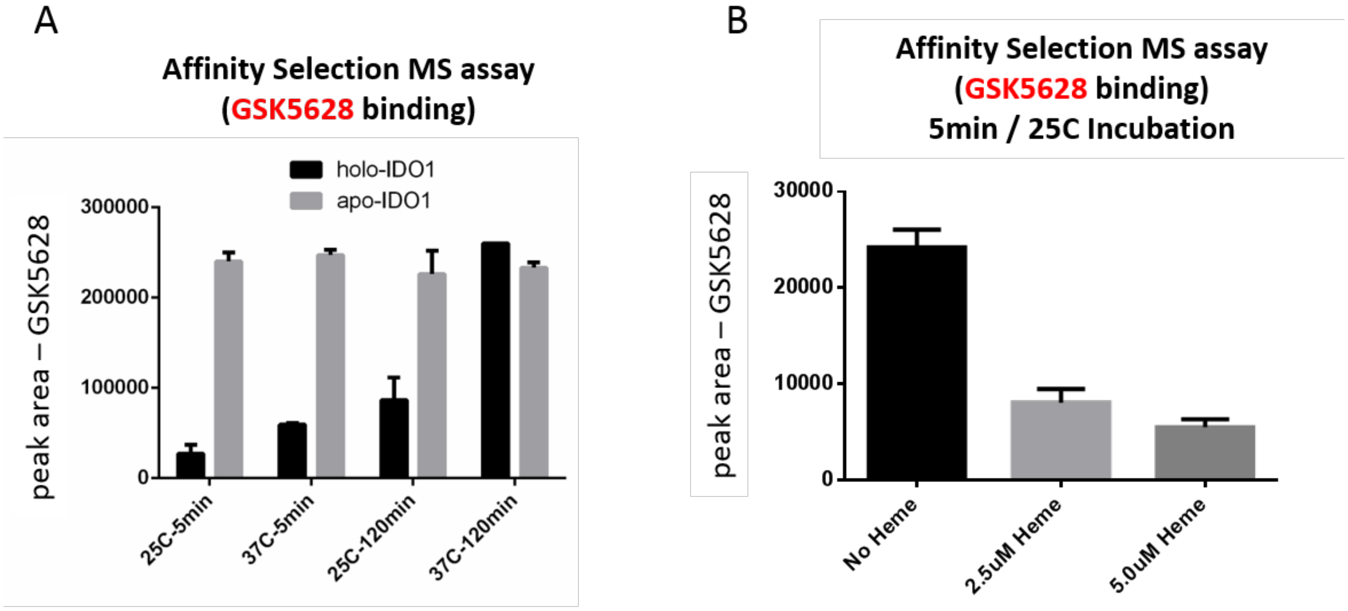


**Figure S12**

Affinity selection / mass spectrometry (AS/MS) experiments to explore GSK5628 binding. (**A**) holo-IDO1 (black bars) or apo-IDO1 (grey bars) were incubated with GSK5628 for the indicated time and the mixture was then passed through a size exclusion column and the amount of GSK5628 was quantitated with an HPLC-MS instrument. Resultant peak area of GSK5628 is represented. These results show that GSK5628 is able to bind apo-IDO1 at all indicated incubation times and temperature but is only able to bind holo-IDO1 as the time and temperature are increased. A control reaction of only GSK5628 showed that passage through the size exclusion column was dependent on the presence of IDO1 (data not shown). (**B**) Apparent binding of GSK5628 to holo-IDO1 (25°C-5 min data) can be reduced by addition of exogenous heme to the incubation. This reduction of binding is consistent with GSK5628 exclusively binding apo-IDO1 (which, even with only a 5 min incubation at 25°C is present in small quantities; note that the y-axis scale is 10 times smaller than **A**), as adding heme to the system pushes the equilibrium toward holo and reduces GSK5628 binding. Results are the average and standard deviation of N=3 experiments.

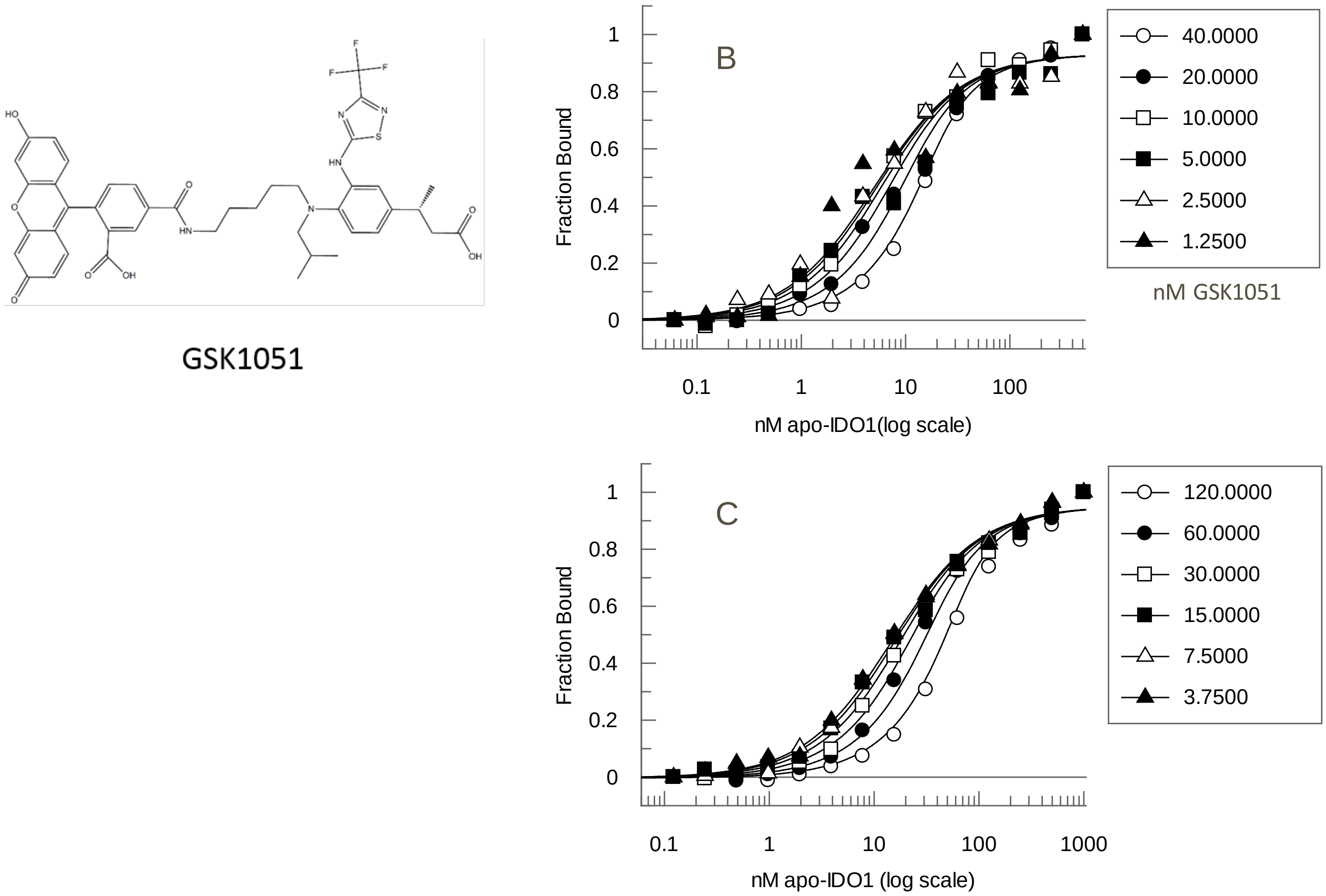


**Figure S13**

(**A**) GSK5628-based fluorescent probe. A fluorescein fluorophore was coupled to a close anlog of GSK5628 with a linker via an NHS-Ester / Amine coupling reaction. The resultant fluorescein-tagged analog (GSK1051) is shown. (**B**) Dual titration of GSK1051 and apo-IDO1 showing that GSK1051 is able to bind apo-IDO1 with an affinity of 10.6 ± 2.46 nM at 25°C. Additional fitted parameters (solid lines) are rfree = -0.0009 ± 0.012, rbound = 0.93 ± 0.013 and a (fractional activity) = 2.3 ± 0.39. (**C**) A similar method was used to measure the probe affinity at 37°C (average Kd = 24.8 ± 7.9 nM from 2 titrations each read at 10, 15 and 20 min; n=6). Fitted parameters for the graph of one measurement at 15 min (solid lines) are Kd = 25.2 ± 2.7 nM, rfree = -0.0035 ± 0.0058, rbound = 0.95 ± 0.0078 and a (fractional activity) = 1.9 ± 0.14.

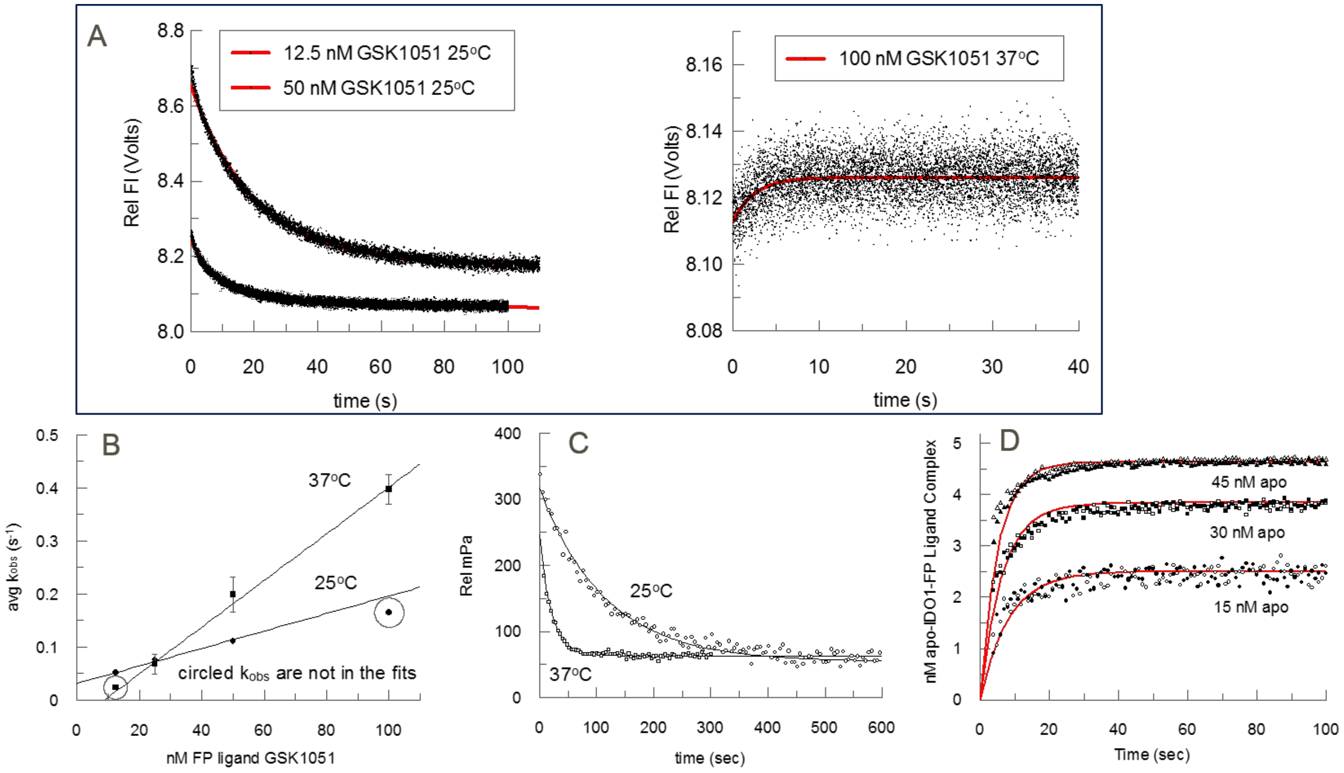


**Figure S14**

Determination of the rate constants for binding of the fluorescent probe GSK1051 to apo-IDO1 at 25°C and 37°C: (**A**) Shows representative average (n = 4 to 5) stopped-flow fluorescence quench time courses for GSK1051 binding to apo-IDO1 fitted to a single exponential (plus a linear region for 25°C) and offset (25°C 12.5 nM avg $k_{obs}$ = 0.0513 ± 0.00017 $s^{-1}$; 25°C 50 nM avg $k_{obs}$ = 0.111 ± 0.000699 $s^{-1}$; 37°C 100 nM avg $k_{obs}$ = 0.397 ± 0.028 $s^{-1}$. In contrast to room temperature the fluorescence increased very slightly at 37°C rather than decreased. (**B**) Shows replots of avg $k_{obs}$ versus GSK1051 to determine $k_{on}$ from the slope (1.65 ± 0.15 $\mu M^{-1}s^{-1}$at 25°C and 4.4 ± 0.35 $\mu M^{-1}s^{-1}$ at 37°C). The intercept at 25°C is ~ $k_{off}$ = 0.031 ± 0.0026 $s^{-1}$, which is within 3-fold of the measured value below, while the intercept at 37°C is poorly defined. (**C**) Shows representative probe $k_{off}$ determinations by competition with an excess of unlabeled GSK5628 (25°C n = 3 avg $k_{off}$ = 0.00884 ± 0.00031 $s^{-1}$ and 37°C n = 6 avg $k_{off}$ = 0.045 ± 0.0025 $s^{-1}$). The calculated probe Kd = $k_{off}/k_{on}$ = 5.4 nM at 25°C and 10 nM at 37°C which are within 2.5-fold of the values of 10.6 and 24.8 nM, respectively, obtained by dual titrations of ligand with apo-IDO1 at both temperatures (**Figures S13B and C**). (**D**) Due to the small change in the intrinsic fluorescence at 37°C , the probe $k_{on}$ (2.78 ± 0.073 $\mu M^{-1}s^{-1}$) and $k_{off}$ (0.069 ± 0.00066 $s^{-1}$) were also measured by KinTek global fits of the FP time courses at 15, 30, 45 and 60 nM probe in duplicate. These values are within 2-fold of the values measured by independent methods illustrated in **Figures S14A, B and C**, and the calculated Kd = $k_{off}/k_{on}$ = 24.9 nM, which is in excellent agreement with the titration Kd (**Figure S13C**).

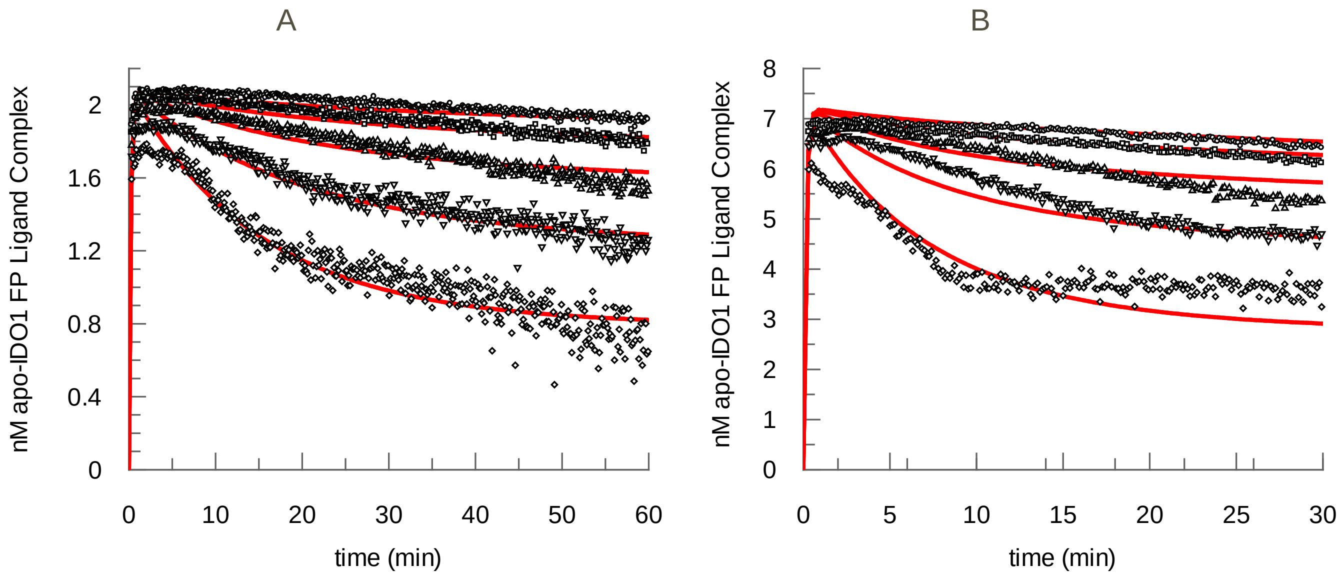


**Figure S15**

Determination of the rate constants for binding of heme by competition with the fluorescent probe GSK1051 at 25°C and 37°C. (**A**) Time course examples with KinTek global fits (solid red lines) for heme competition at 25°C; k1 = 3.4 ± 0.17 $\mu M^{-1}s^{-1}$, k2 = 0.037 ± 0.0019 $s^{-1}$, k3 = 0.067 ± 0.0024 $\mu M^{-1}s^{-1}$ , k4 = 0.00027 ± 0.0000078 $s^{-1}$, k5 = 0.017 ± 0.0029 $s^{-1}$, k6 = 0 (locked), where k1 and k2 are probe $k_{on}$ and $k_{off}$, k3 and k4 are heme $k_{on}$ and $k_{off}$, and k5 is the signal decay rate. For clarity curves are shown for only one duplicate at 1.25 to 20 nM heme in 2-fold increments (top to bottom); global fits are to 0, 0.3125 to 40 nM heme in duplicate. The calculated Kd values from $k_{off}/k_{on}$ = 11 nM for the probe and 4 nM for heme at 25°C. (**B**) Time course examples with KinTek global fits (solid red lines) for heme competition at 37°C; 2.2 ± 0.13 $\mu M^{-1}s^{-1}$, k2 = 0.055 ± 0.0032 $s^{-1}$, k3 = 0.098 ± 0.0039 $\mu M^{-1}s^{-1}$, k4 = 0.00073 ± 0.000026 $s^{-1}$, k5 = 0.041 ± 0.0041 $s^{-1}$, k6 = 0 (locked). For clarity curves are shown for only one duplicate at 2.5 to 40 nM heme in 2-fold increments increments (top to bottom); global fits are to 0, 0.3125 to 40 nM heme in duplicate. The calculated Kd values from $k_{off}/k_{on}$ = 25 nM for the probe and 7.5 nM for heme at 37°C. The heme (k3 and k4) and probe (k1 and k2) rate constants compare favorably (within 4-fold) with those measured independently in **Figure 4B and S13** respectively.

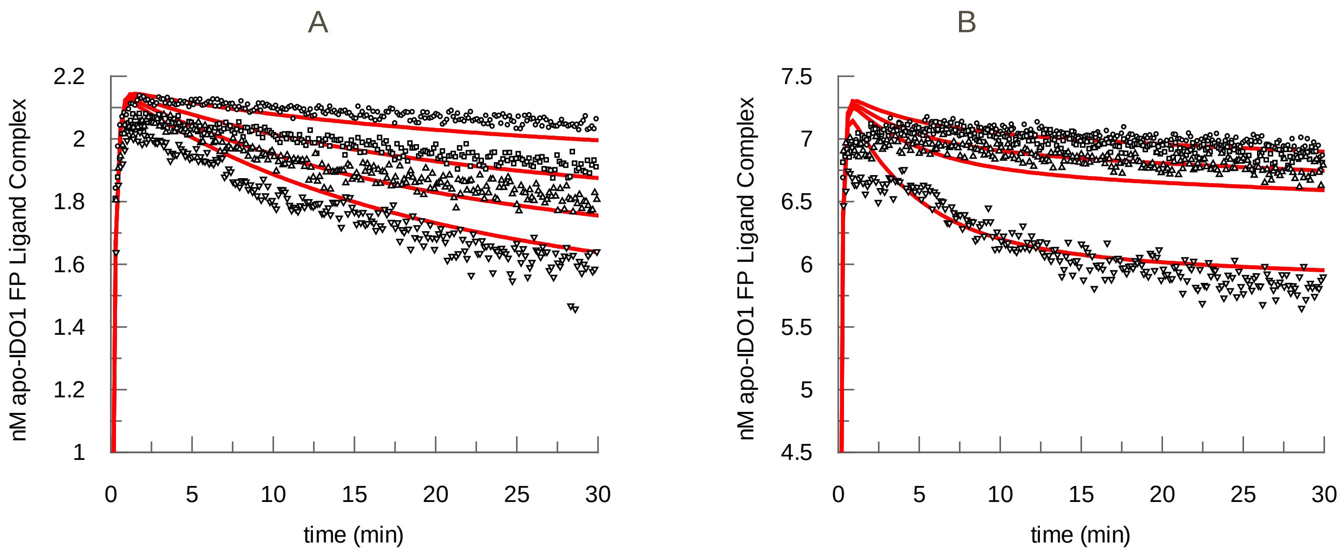


**Figure S16**

Determination of the rate constants for binding of GSK5628 by competition with the fluorescent probe GSK1051 at 25°C and 37°C. (**A**) Time course examples with KinTek global fits (solid red lines) for GSK5628 competition at 25°C; k1 = 3.7 ± 0.16 $\mu M^{-1}s^{-1}$, k2 = 0.039 ± 0.0017 $s^{-1}$, k3 = 0.11± 0.0072 $\mu M^{-1}s^{-1}$, k4 = 0.00015 ± 0.000020 $s^{-1}$, k5 = 0.023 ± 0.0033 $s^{-1}$ and k6 = 0 (locked), where k1 and k2 are probe $k_{on}$ and $k_{off}$, k3 and k4 are GSK5628 $k_{on}$ and $k_{off}$, and k5 is the signal decay rate. For clarity curves are shown for only one duplicate at 1.25, 2.5, 3.75 and 5 nM GSK5628 (top to bottom); global fits are to 0, 1.25, 2.5, 3.75 and 5 nM GSK5628 in duplicate. The calculated Kd values from $k_{off}/k_{on}$ = 10 nM for the probe and 1.3 nM for GSK5628 at 25°C. (**B**) Time course examples with KinTek global fits (solid red lines) for GSK5628 competition at 37°C; k1 = 2.0 ± 0.10 $\mu M^{-1}s^{-1}$, k2 = 0.047 ± 0.0024 $s^{-1}$, k3 = 0.15 ± 0.016 $\mu M^{-1}s^{-1}$ , k4 = 0.00064 ± 0.000053 $s^{-1}$, k5 = 0.025 ± 0.0019 $s^{-1}$ and k6 = 0 (locked), where k1 and k2 are probe $k_{on}$ and $k_{off}$, k3 and k4 are GSK5628 $k_{on}$ and $k_{off}$, and k5 is the signal decay rate. For clarity curves are shown for only one duplicate at 2.5, 3.75, 5 and 10 nM GSK5628 (top to bottom); global fits are to 0, 1.25, 2.5, 3.75, 5, 7.5 and 10 nM GSK5628 in duplicate. The calculated Kd values from $k_{off}/k_{on}$ = 23 nM for the probe and 4.2 nM for GSK5628 at 37°C. The GSK5628 and probe rate constants compare favorably (within 4-fold) with those measured independently in **Figures S10** (slow phase)**, S11** (GSK5628) **and S14** (probe), respectively**.**

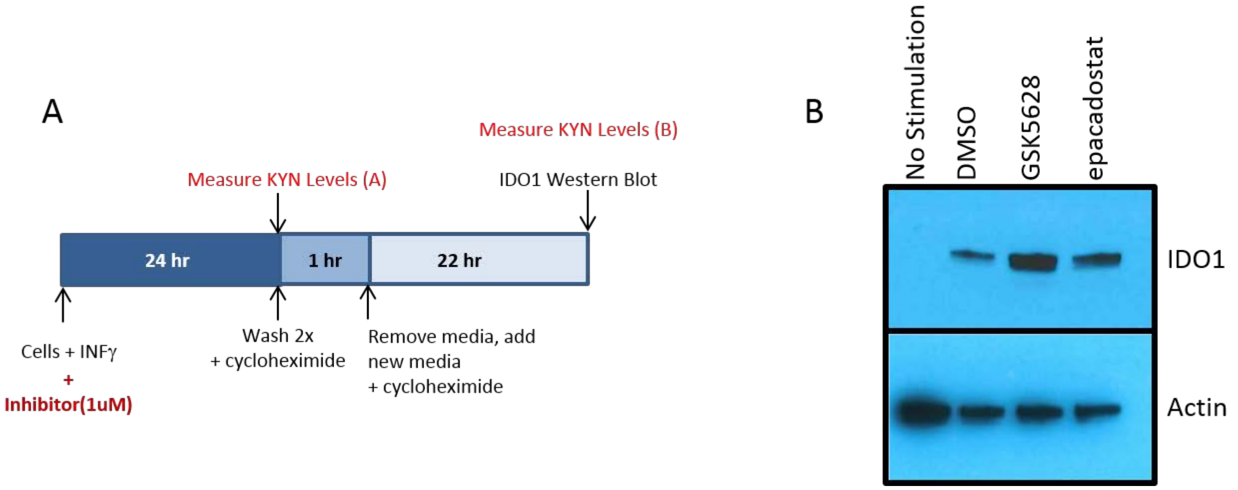


**Figure S17**

GSK5628 washout experiments on IFN-γ stimulated HeLa cells. (**A**) Scheme of the experiment shown in **Figure 5A**: a wash-out combined with KYN levels measurements to assess IDO1 activity. (**B**) Cell lysates were prepared from HeLa cells at the indicated time point and were subjected to SDS-PAGE, transferred to a nitrocellulose membrane and probed with the indicated antibody. Results show that IDO1 (Antibody: Cell signaling 86630) is not degraded upon treatment with both GSK5628 and epacadostat. Actin (Antibody: Cell signaling 3700) is present as a loading control.

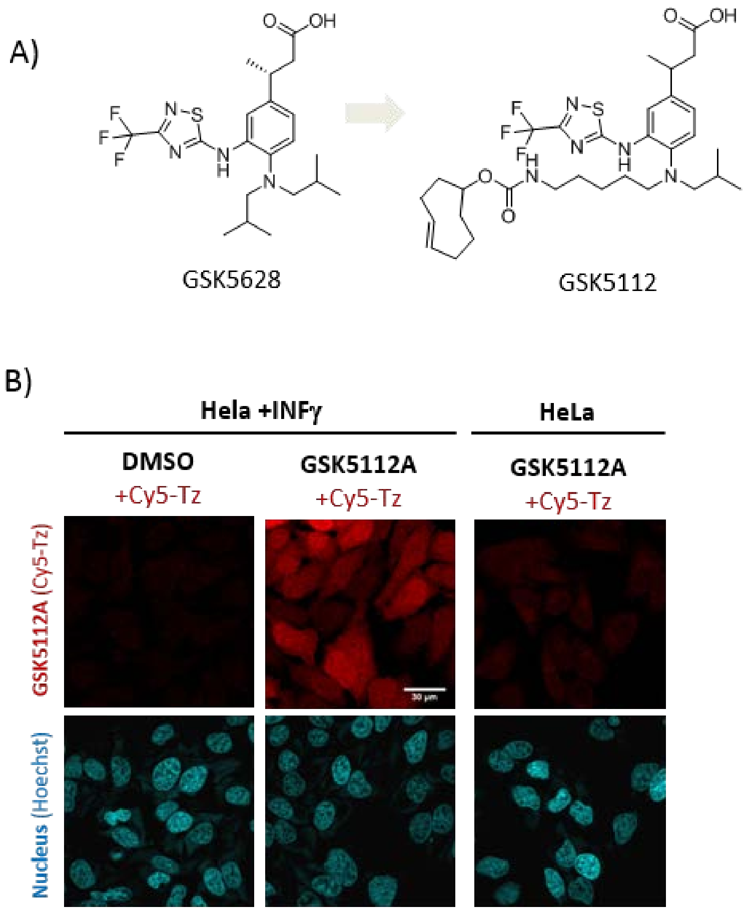


**Figure S18**

(**A**) Click-probe GSK5112 derived from novel IDO1 inhibitor GSK5628; (**B**) Representative fluorescent images of none-stimulated and INFγ stimulated HeLa cells, which were treated with click-probe GSK5112 (0.15 µM) for 60 min followed by fixation, permeabilization and click reaction with 100 nM Cy5-Tz for 5 min as well as by Hoechst staining. Images were recorded after excitation at 633 nm (Cy5, upper panel) and at 405 nm (Hoechst, lower panel). Scale bar = 30 µm.

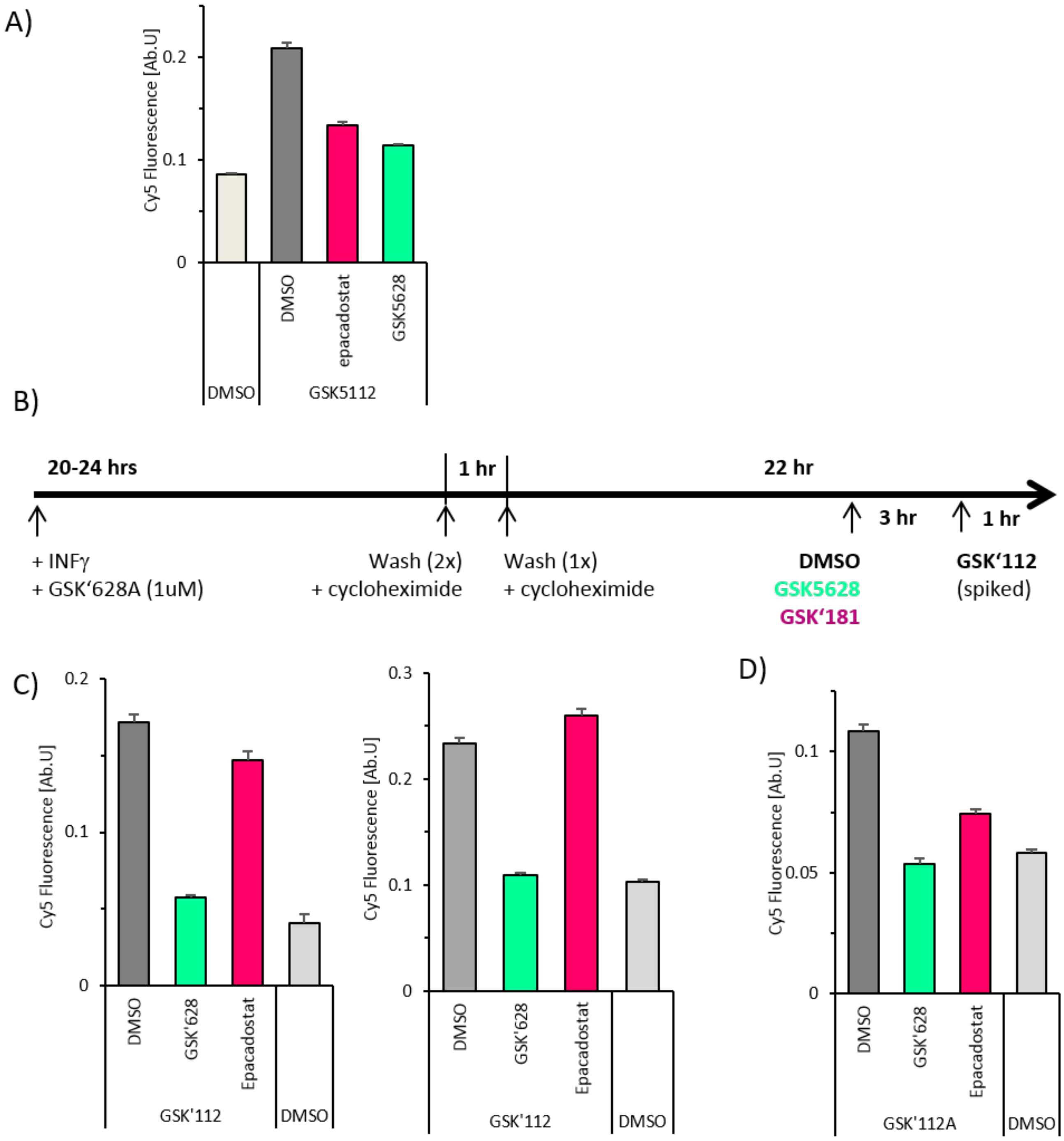
A)
Cy5 Fluorescence [Ab.U]
0.2
0.1
0
DMSO
epacadostat
GSK5628
DMSO
GSK5112
B)
20-24 hrs
1 hr
22 hr
3 hr
1 hr
+ INFγ
+ GSK'628A (1uM)
Wash (2x)
+ cycloheximide
Wash (1x)
+ cycloheximide
DMSO
GSK5628
GSK'181
GSK'112
(spiked)
C)
Cy5 Fluorescence [Ab.U]
0.2
0.1
0
DMSO
GSK'628
Epacadostat
GSK'112
DMSO
Cy5 Fluorescence [Ab.U]
0.3
0.2
0.1
0
DMSO
GSK'628
Epacadostat
GSK'112
DMSO
D)
Cy5 Fluorescence [Ab.U]
0.1
0.05
0
DMSO
GSK'628
Epacadostat
GSK'112A
DMSO

**Figure S19**

(**A**) Quantification of a representative competition experiment shown in **Figure 5B**. Mean fluorescence (Cy5 channel) of individual INFγ stimulated HeLa cells (n=93-119) has been quantified from 6 representative images of cells treated with either DMSO, epacadostat or GSK5628 prior to the addition of click-probe GSK5112. SEM is shown. (**B**) Scheme of the experiment shown in **Figure 5C**: a wash-out combined with click-probe treatment. (**C**) Quantification of two additional, independent, experiments described in **Figure 5C**. Mean fluorescence (Cy5 channel) of individual INFγ stimulated HeLa cells (n≥90) has been quantified from at least 5 representative images of cells. SEM is shown. (**D**) Quantification of a control wash-out experiment to test if cycloheximide treatment and wash steps can influence the IDO1 activity per se. Experiment was performed as described in **Figure S19B** but without addition of GSK5628 at time point zero. SEM is shown.

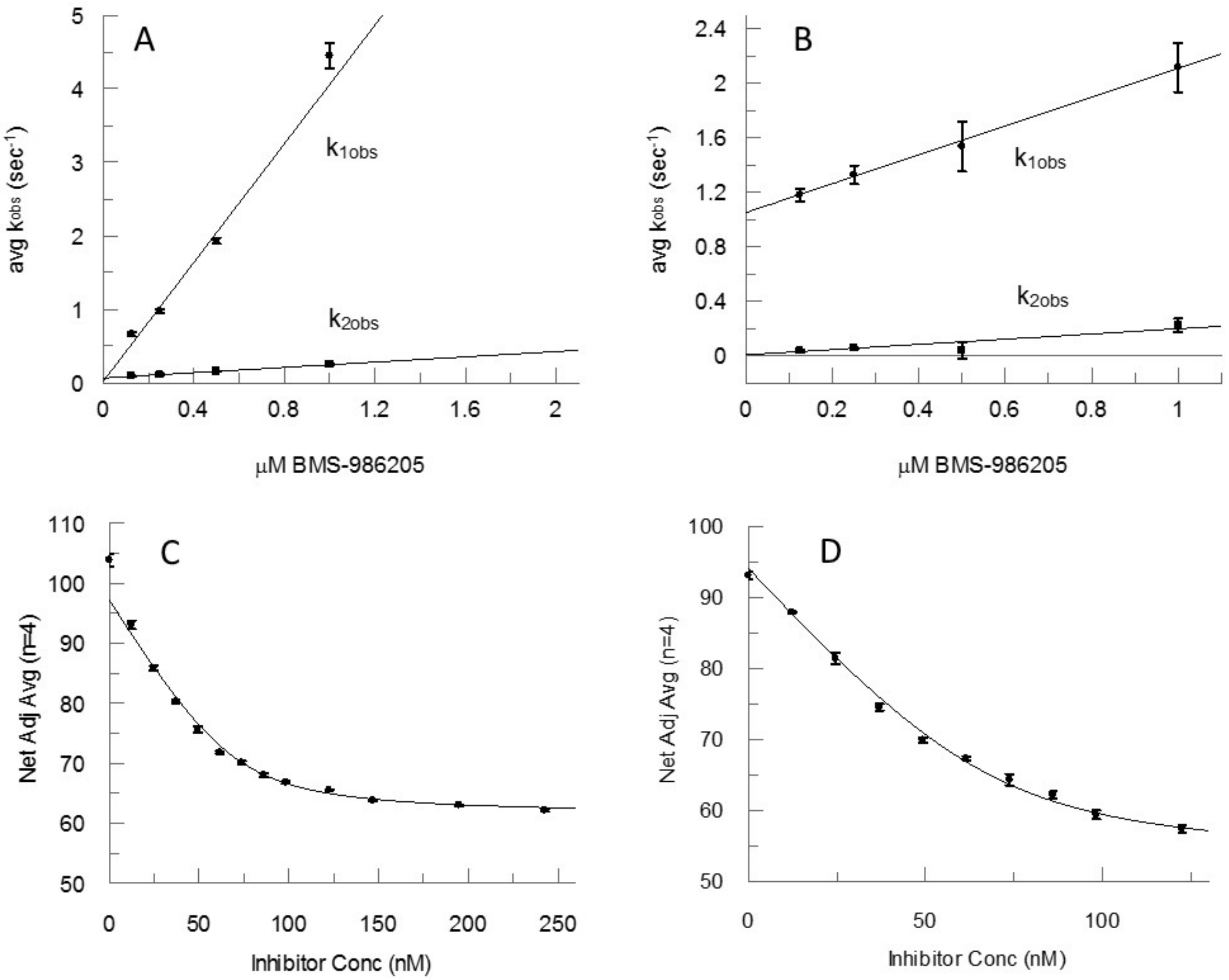


**Figure S20**

Stopped-flow and equilibrium titrations of the quenching of intrinsic apo-IDO1 trp fluorescence with BMS-986205 (BioVision Inc.) at 25°C and 37°C to determine kon and Kd at both temperatures under conditions described in **Figures S10** and **S11**. Time courses (avg n=3) of the quenching of intrinsic apo-IDO1 (50 nM) trp fluorescence were fitted to double exponentials and linear replots of $k_{1obs}$ and $k_{2obs}$ versus BMS-986205 concentrations at 23°C (**A**) and 37°C (**B** ) were used to determine $k_{on}$ from the replot slopes and to demonstrate that direct binding to apo-IDO1 is rapid regardless of temperature. Replots are linear fits of the averages of n=3 runs and the standard deviations using GraFit (Erithacus Software). Values for $k_{on}$ (slopes) are 4.0 ± 0.48 $\mu M^{-1}s^{-1}$ for $k_{1obs}$ (r = 0.980) and 0.18 ± 0.0097 $\mu M^{-1}s^{-1}$ for $k_{2obs}$ (r = 0.997) at 23°C and 1.1 ± 0.043 $\mu M^{-1}s^{-1}$ for $k_{1obs}$ (r = 0.998) and 0.19 ± 0.043 $\mu M^{-1}s^{-1}$ for $k_{2obs}$ (r = 0.953) at 37°C. Intercepts poorly define $k_{off}$ due to extreme extrapolation from the lowest concentrations to the Kd's.

Equilibrium titrations of the quenching of intrinsic apo-IDO1 trp fluorescence with BMS-986205 at 25°C (**C**) and 37°C (**D**) were used to determine an accurate Kd at both temperatures. Fitted parameters shown by the solid lines are (**C**): Kd = 6.9 ± 1.4 nM, E locked at 70 nM, A = 97 ± 1.2 RFU and B = 61 ± 0.56 RFU; (**D**): Kd = 7.9 ± 4.5 nM, E locked at 70 nM, A = 94 ± 0.43 RFU and B = 53 ± 3.6 RFU.

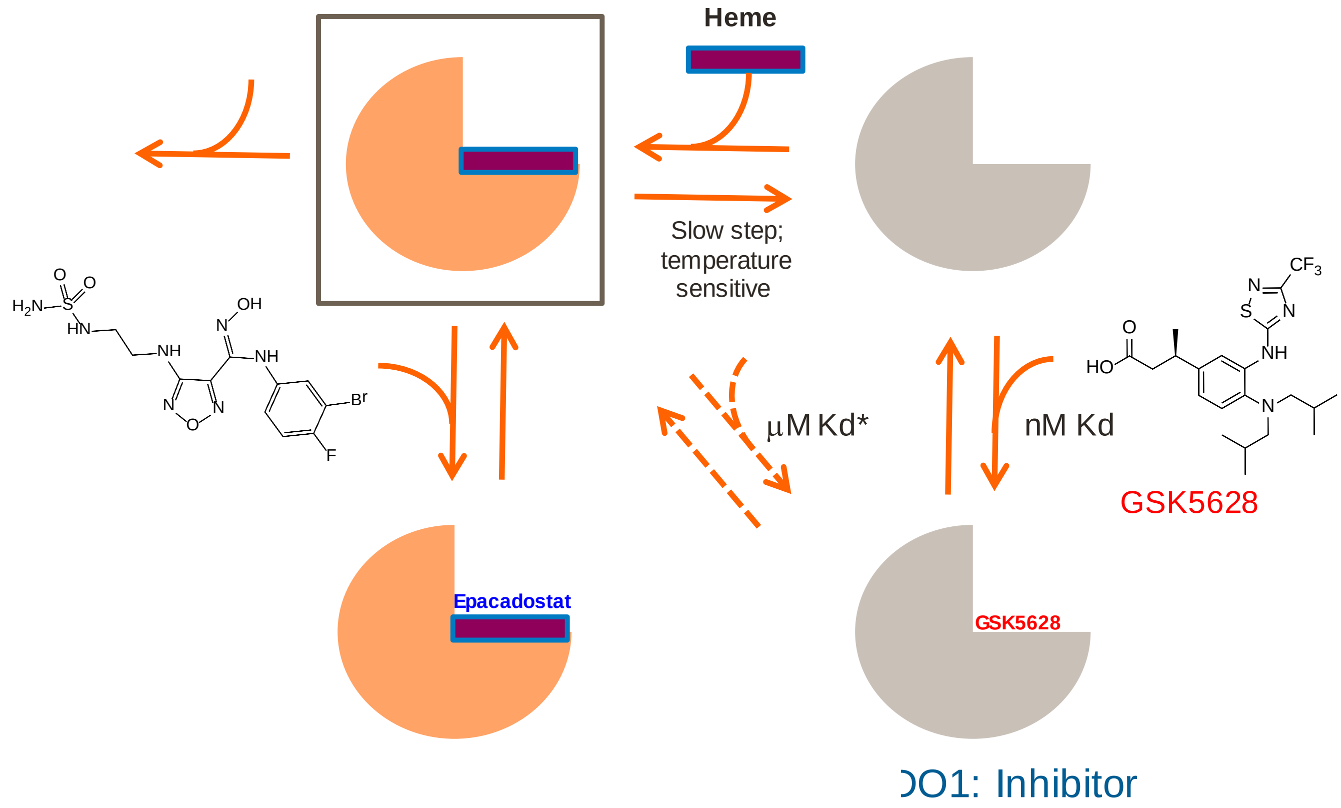


**Figure S21**

Scheme of equilibrium between holo-IDO1 and apo-IDO1 depicting different binding modes of epacadostat and GSK5628 including a low affinity pathway for compounds related to GSK5628 and BMS-986205. The low affinity pathway acknowledges the crystal structures identified by Pham and Yeh, 2018.
